# Cell Diameter in *Bacillus subtilis* is Determined by the Opposing Actions of Two Distinct Cell Wall Synthetic Systems

**DOI:** 10.1101/392837

**Authors:** Michael F. Dion, Mrinal Kapoor, Yingjie Sun, Sean Wilson, Joel Ryan, Antoine Vigouroux, Sven van Teeffelen, Rudolf Oldenbourg, Ethan C. Garner

## Abstract

Rod shaped bacteria grow by adding material into their cell wall via the action of two spatially distinct enzymatic systems: The Rod system moves around the cell circumference, while the class A penicillin-binding proteins (aPBPs) are unorganized. To understand how the combined action of these two systems defines bacterial dimensions, we examined how each system affects the growth and width of *Bacillus subtilis*, as well as the mechanical anisotropy and orientation of material within their sacculi. We find that rod diameter is not determined by MreB, rather it depends on the balance between the systems: The Rod system reduces diameter, while aPBPs increase it. RodA/PBP2A can both thin or widen cells, depending on its levels relative to MreBCD. Increased Rod system activity correlates with an increased density of directional MreB filaments, and a greater fraction of directionally moving PBP2A molecules. This increased circumferential synthesis increases the amount of oriented material within the sacculi, increasing their mechanical anisotropy and reinforcing rod shape. Together, these experiments explain how the combined action of the two main cell wall synthetic systems build rods of different widths, a model that appears generalizable: *Escherichia coli* containing Rod system mutants show the same relationship between the density of directionally moving MreB filaments and cell width.

## Introduction

While the length of *Bacillus subtilis* rods increases as a function of their growth rate ^1^, their width remains constant across different growth conditions ^2^. How bacteria define and maintain these rod shapes with such precision is not understood, but it must involve mechanisms controlling both the rate and location of glycan insertion into the peptidoglycan (PG) sacculus, the enveloping heteropolymer meshwork that holds cells in shape ^3^. In order to understand how bacteria grow in defined shapes, we need to understand not only where these enzymes act, but how their activity affects the geometry and arrangement of material within the sacculus.

The PG for cell elongation is synthesized by two families of penicillin-binding proteins (PBPs). The class A penicillin-binding proteins (aPBPs) both polymerize and cross-link glycans. The class B penicillin-binding proteins (bPBPs) cross-link ^4, 5^ the glycans polymerized by the glycosyltransferase RodA ^6^. bPBPs and RodA are components of the “Rod complex”, a group of proteins essential for rod shape **(Figure 1A).** In *B. subtilis*, this contains RodA, the class B transpeptidase PBP2A, MreC, MreD, RodZ, and filaments of MreB, an actin homolog ^7, 8^. MreB polymerizes together with other MreB homologs into short, highly curved filaments on the membrane ^9, 10^. To maximize their interaction with the membrane these curved filaments deform to the membrane, orienting along the direction of highest inward membrane curvature, pointing around the rod width^11^. Oriented by their association with MreB filaments, the Rod complex moves directionally around the cell circumference, driven by the PG synthetic activity of RodA/PBP2A ^12^-^15^. This radial motion of independent filament-enzyme complexes are believed to insert PG strands around the rod width ^16^.

**Figure 1.**
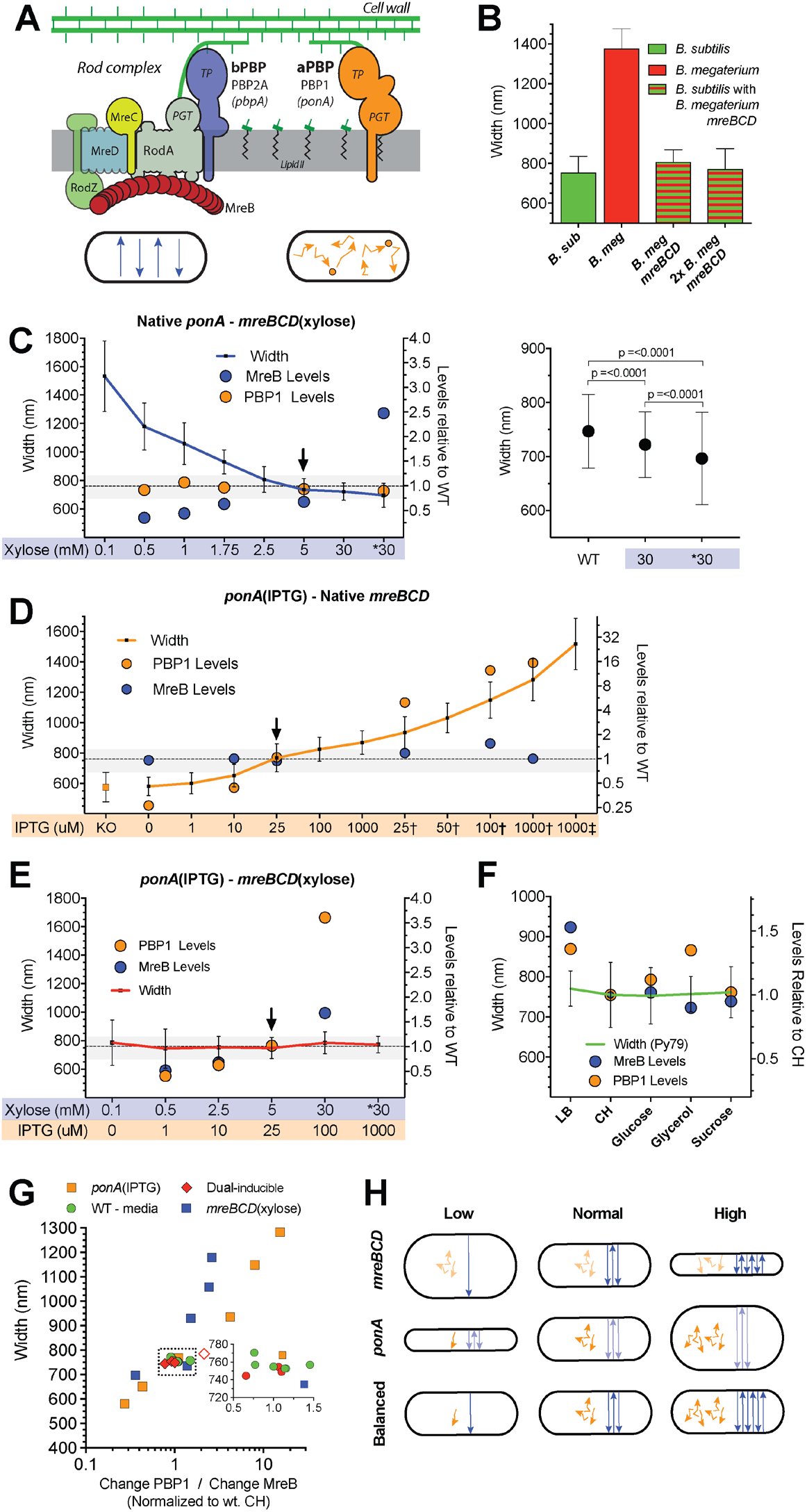
Rod width depends on the relative levels of the widening aPBPs to the thinning Rod system. Error bars are SD of the mean. **A. Schematic of the two PG synthetic systems responsible for growth**. The *Rod complex* (depicted on the left) polymerizes glycans via the peptidoglycan glycosyltransferase (PGT) activity of RodA. These glycans are crosslinked into the sacculus by the transpeptidase (TP) activity of PBP2A. *aPBPs* are depicted on the right, containing both PGT and TP activity. *Below* - Schematic of each system’s *in vivo* motions. Rod complexes move circumferentially around the rod as they synthesize PG. In contrast, aPBP synthesis does not appear organized, as they diffuse along the membrane, occasionally halting and remaining immobile for 1-3 seconds. Protein names are capitalized, gene names italicized. **B. *B. megaterium mreBCD* in *B. subtilis* forms rods close to *B. subtilis* width.** “*B. sub”* indicates WT *B. subtilis.* “*B. meg****”*** indicates *B. megaterium*. Checkered boxes are bMD465 *(amyE::Pxyl-mreBCD, minCD [B. mega]::erm, mreBCD, minCD::spec [B. mega])*, a *B. subtilis* strain where the native *mreBCD* locus is replaced with *B. megaterium mreBCD*, and an additional *B. megaterium mreBCD* is under xylose control at an ectopic locus. “*B. meg mreBCD*” indicates bMD465 grown with 1% glucose to repress expression from the additional copy. “*2x B. meg mreBCD”* indicates bM465 growth with 30 mM xylose to overexpress *megaterium mreBCD* from the additional copy. See **Figure S1A** for a zoomed view. **C-F. Titrations of *ponA* and *mreBCD* vs. cell width.** Each strain was grown in the inducer concentrations shown below the graph. Mean width is plotted on the left axis. PBP1 and MreB relative protein abundances (determined by mass spectrometry and normalized to the levels in WT PY79 grown in CH) are plotted on the right axis. Arrowhead indicates inductions producing WT widths and cellular levels of induced protein. Dashed line depicts mean steady state diameter of PY79 in CH, with SD shown as gray shaded area. **C. Cell diameter decreases with *mreBCD* induction**. All data are from strain bMD545 (*amyE::Pxyl-mreBCD::erm ΔmreBCD::spec)*, except for the induction marked **\*** which are bMK355 (*amyE::Pxyl-mreBCD::erm*) containing a xylose-inducible *mreBCD* in addition to the native *mreBCD*. *Right –* Zoomed view of the right two bars compared to WT. P-value calculated with Mann-Whitney. **D. Cell diameter increases with *ponA* induction**. All data are from strain bMD598 (*yhdG::Pspank-ponA*::*cat* Δ*ponA::kan*), except for the inductions marked † and ‡ which are under stronger promoters than bMD598. † is bMD586 (*yhdG:Phyperspank*-*ponA*::*cat* Δ*ponA::kan*), and ‡ is bMD554 (*yhdG::Phyperspank-ponA*::*cat*) which has an inducible *ponA* in addition to the native copy. **E. A balanced expression of both PG synthetic systems yields normal width rods across a large range.** Strain bMD620 (*amyE::Pxyl-mreBCD*::*erm* Δ*mreBCD::spec yhdH::Pspank*-*ponA*::*cat* Δ*ponA::kan*) grown in combinations of inducers shown along the bottom. **\*** indicates bMD622 (*amyE::Pxyl-mreBCD*::*erm yhdH::Pspank*-*ponA*::*cat* Δ*ponA::kan*) containing a xylose-inducible *mreBCD* in addition to the native *mreBCD.* **F. *B. Subtilis* maintains a constant width in different media.** Mean and SD of cell diameters of *B. subtilis* strain PY79 (exponential phase) grown in different growth media (media where a carbon source is indicated is S750). Width is plotted on the left axis. PBP1 and MreB relative protein abundances (determined by mass spectrometry, normalized to the levels in CH media) are plotted on the right axis. **G. WT width is maintained within a narrow range of relative PBP1/MreB ratios.** Plotted are the widths of all cells in Figures 1C-F, against the ratio of the fold change in PBP1 to that of MreB. Inset is a zoomed view of the dotted box, showing the widths of cells within a 0.5-1.5 PBP1/MreB range. Green indicates ratios in different media. Orange squares indicate ratios in *ponA* inducible strains. Blue squares indicate ratios in *mreBCD* inducible strains. Red diamonds indicate ratios in the dual-induction strain, with the open red diamond indicating the ratio when protein levels exceeded WT. **H. Model for how the two PG synthesis systems affect rod width.** *Top* – As the amount of circumferentially organized synthesis increases (blue straight lines), cell diameter decreases. *Center* – In contrast, as the level of unorganized synthesis increases (orange, non-directed lines), so too does cell diameter. *Bottom* – As long as the activities of the unorganized (orange) and circumferentially organized (blue) syntheses are balanced, cell width can remain constant, even across a range of protein levels.

aPBPs also affect rod shape, as *B. subtilis* cells lacking them are thinner ^17^. Single molecule and biochemical studies indicate aPBPs and the Rod system are both spatially and functionally distinct ^6, 14^: While Rod complexes move around the cell width, aPBPs have never been seen to move directionally. Rather, they diffuse within the membrane, occasionally pausing and remaining immobile for a few seconds ^14^. Furthermore, the synthetic activity of either aPBPs or the Rod complex can be inhibited without affecting the activity or motions of the other ^6, 14^. Given that both synthetic systems are required for growth, but that Rod complex activity is spatially organized while aPBP activity is not **(Figure 1A)**, it is not clear how these two PG synthetic machineries work together to create a rod-shaped sacculus of a given width.

Current models of rod width have focused on MreB filaments, attributing the altered widths of *Escherichia coli* MreB mutants to possible changes in MreB filament curvature, twist, angle, or localization to negative Gaussian curvature ^18^-^24^. Not only do these models lack the effects of aPBP mediated synthesis, they are: A) theoretical, as these proposed changes in filament curvature or twist have not been structurally validated, or B) difficult to reconcile in *B. subtilis*, which has no detectable negative Gaussian curvature ^11^ or skew in filament angle relative to the cell body ^11^-^13, 25^.

Rather than focusing on MreB alone, we sought to develop a more thorough understanding of how the synthesis of both organized and unorganized PG affects the width and growth of *B. subtilis*, as well as their effects on the organization and mechanics of cell wall material. We find that aPBP and Rod complex-mediated PG synthesis have opposing effects on rod width, and that cell diameter depends on their balance. The rate cells add surface area is largely unaffected by the level of any one system, unless both become limiting. As MreBCD expression increases and rods thin, we observe a greater density of directionally moving MreB filaments and a greater fraction of directionally moving enzymes. Increased Rod complex activity creates a greater proportion of oriented material within the sacculus, causing the rod to stretch less across its width and more along its length in response to internal turgor. Finally, we show that the different widths of *E. coli* Rod mutants also show correlation between the density of directionally moving MreB filaments and cell width, giving a simple, generalizable model that may explain the role of the Rod system in cell width.

## Results

### The Rod and aPBP systems have opposing effects on cell diameter

The width of rod-shaped bacteria has been attributed to properties encoded within MreB filaments ^19, 21, 24, 26^. We reasoned that if width were defined by MreB, then the *mreBCD* genes from larger bacteria should produce cells with larger diameters. To test this, we replaced the *B. subtilis mreBCD* operon with the *mreBCD* operon from *Bacillus megaterium*, a nearly 2-fold wider bacterium. Surprisingly, these cells grew as rods only slightly wider than wild type (WT) *B. subtilis*, and by further overexpressing *B. megaterium mreBCD* cells became even thinner (36 nm) **(Figure 1B, S1A)**, suggesting that MreB filaments themselves do not encode a specific rod width.

We next examined how width changed as we independently controlled the levels of the two main *B. subtilis* PG synthetic systems. We created strains where expression of PBP1 (the major aPBP, encoded by *ponA*) was under IPTG control. As PG synthesis by the Rod system depends upon MreB ^14^, we created strains where the native *B. subtilis mreBCD* operon was under xylose control. We grew these strains at different induction levels, then measured their steady-state widths using fluorescence microscopy. For a subset of inductions, we measured the relative protein abundance using proteomic mass spectrometry, normalizing to the levels in WT cells grown in CH media (**Supplemental File 1**).

Varying *mreBCD* inductions revealed the Rod system has a thinning effect on rod shape. In the lowest inductions supporting growth, rods were ∼2-fold wider than WT, with some cells losing rod shape. As we increased *mreBCD* induction, cells became thinner, reaching WT width when MreB abundance recapitulated WT levels **(Figure 1C, Figure S1B).** Inductions above this point resulted in cells becoming 33 nm thinner than WT, and thinner still (58 nm) in *mreBCD* merodiploids. As above, these results demonstrate MreB filaments do not *define* a given rod diameter, rather as previously hypothesized ^11, 27^, they indicate the Rod system *reduces* cell diameter.

Different *ponA* inductions revealed that aPBPs have a widening effect: With no IPTG, cells contained 0.25 the amount of WT PBP1, and were ∼23% thinner than WT, similar to Δ*ponA* strains ^17^ **(Figure 1D, Figure S1C).** As we increased *ponA* induction cells became wider, reaching WT widths when PBP1 abundance recapitulated WT levels. Inductions above this point caused cells to become increasingly wider, and by expressing *ponA* under stronger promoters or in merodiploids, we could produce rods almost twice WT diameter.

These results demonstrate that the aPBPs and Rod system have opposing effects: The circumferentially moving Rod complex reduces cell diameter, while the spatially unorganized aPBPs increase diameter. We hypothesized that a balanced expression of each system could produce WT diameter rods. We combined the IPTG-inducible *ponA* and xylose-inducible *mreBCD* alleles into one “dual-inducible” strain. We found six different pairs of xylose and IPTG concentrations that produced rods of WT diameter **(Figure 1E, Figure S1D)**, even though individually, each induction resulted in perturbed diameters in the singly-inducible parental strains **(Figure 1C-D).** Relative quantitation of protein levels revealed that, in the induction pairs at or beneath WT levels, these cells contained reduced, but relatively balanced amounts of PBP1 relative to MreB. To determine this balance in native conditions, we measured the widths and protein levels of WT cells grown in different media that result in different growth rates **(Figure 1F, Figure S1E).** Together, this data suggested width depends on the balance between the aPBP and the Rod systems. By plotting the ratio of the [Fold change PBP1] / [Fold change MreB] for all conditions in the data sets against their width **(Figure 1G)**, we found that *B. subtilis* maintains its diameter within ∼5% of WT only when this ratio was within a range of 0.8 to 1.5; outside of this range, cell diameter diverged. WT cells in different media showed PBP1/MreB ratios ranging from 0.9 to 1.5. Likewise, the induction pairs yielding WT widths in the dual-inducible strain were within 0.8 to 1.0, only when their levels did not exceed WT. In contrast, the single-inducible *ponA* or *mreBCD* strains were within 5% of WT width only when the ratio was within this range; outside of it, width rapidly diverged. Together, these results indicate that, at the level of PG insertion, cell width is affected by the levels of the two opposing synthetic systems that insert material into the sacculus **(Figure 1H).**

**RodA can function outside of the Rod system to widen cells, but only when PBP2A is also in excess.** Given that RodA acts within the Rod complex with PBP2A (encoded by *pbpA*) to synthesize PG ^14^, titrations of *rodA* and *pbpA* should show the same “thinning effect” as *mreBCD*. However, *rodA* overexpression can restore WT width to ΔaPBP cells ^6^, a widening activity similar to *ponA.* To investigate this discrepancy, we made two sets of strains where *rodA* or *pbpA* (as the only elongation-specific bPBP in the cell) were under the control of increasingly strong inducible promoters. As before ^28, 29^, low *rodA* or *pbpA* yielded wide cells; and as induction increased, cells gradually thinned to WT width, suggesting both are required for functional Rod complexes. However, while we expected excess RodA to create thinner cells, once *rodA* induction exceeded the level required for normal width, it behaved like excess *ponA*, fattening cells far beyond WT diameter ^6^ **(Figure 2A).** In contrast, *pbpA* inductions beyond this point had a negligible effect on diameter.

**Figure 2.**
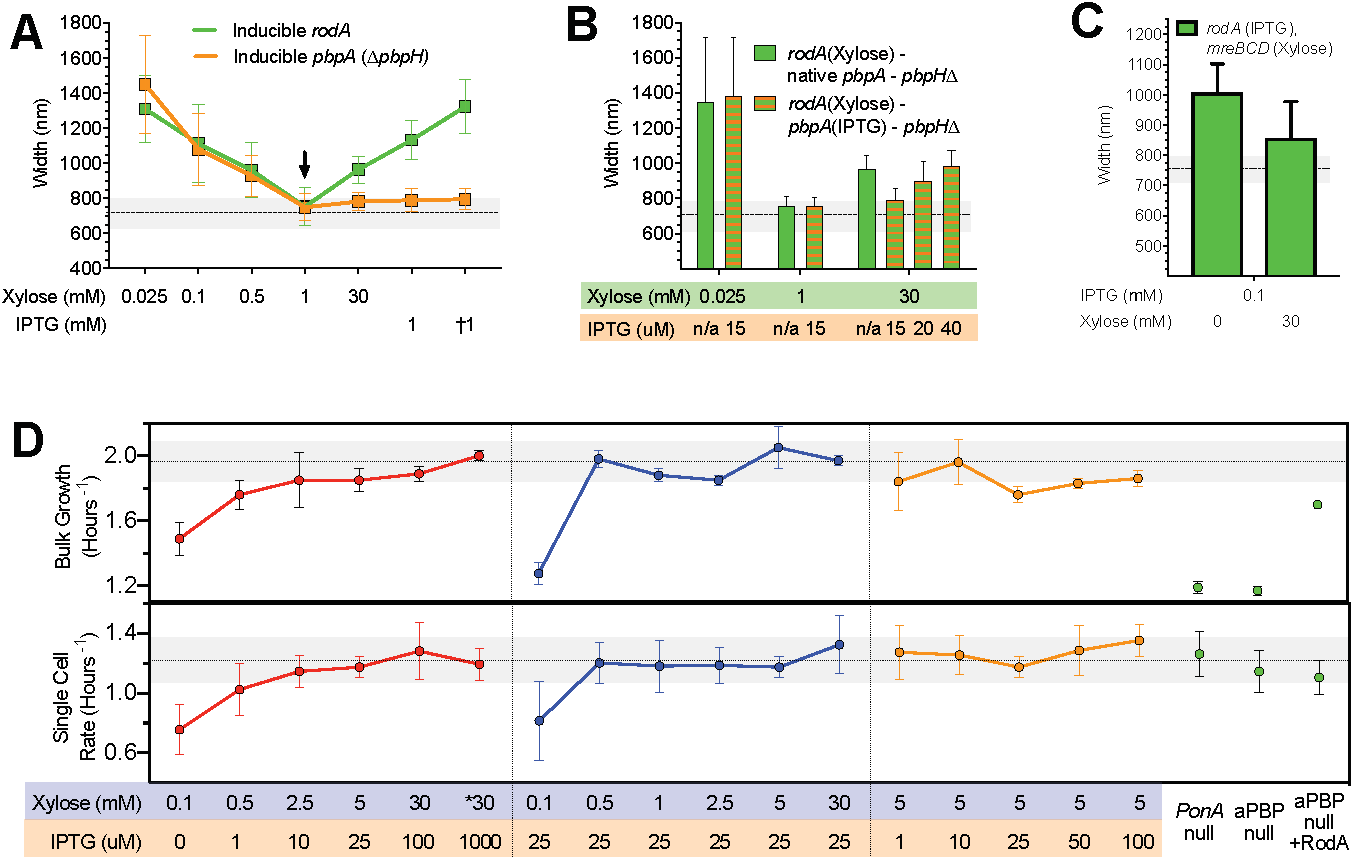
Effects of RodA/PBP2 on cell width, and how each PG synthetic system affects growth. **A. As *rodA* or *pbpA* induction is increased, the mean diameter of cells decreases up to a point, beyond which it increases with rising *rodA* induction, but not for *pbpA*.** Green line represents diameters of bMD592 (Pxyl-*rodA*::*erm*) at five different xylose concentrations, save bMD580 (*yhdG*::Phyperspank-*rodA*::*cat* Δ*rodA::kan*) and bMD556 (*yhdG*::Phyperspank-*rodA*::*cat* - labeled †) which were induced with IPTG. Orange line represents the diameters of bMD597 (Pxyl-*pbpA*::*erm* Δ*pbpH::spec*) at five different xylose concentrations, save bMD574 (*yhdG*::Phyperspank-*pbpA*::*cat* Δ*pbpA::erm* Δ*pbpH::spec*) and bMD573 (*yhdG*::Phyperspank-*pbpA*::*cat* Δ*pbpH::spec* - labeled †) which were induced with IPTG. **B. Overexpression of *rodA* increases cell diameter, but only when *pbpA* expression is also sufficiently high.** Light green bars represent the steady state diameter of bMD627 (Pxyl-*rodA*::*erm* Δ*pbpH::spec*) induced at the specified concentrations of xylose. Green + orange striped bars represent the steady state diameter of bMD631 (Pxyl-*rodA*::*erm yhdG*::Pspank-*pbpA::phleo* Δ*pbpH::spec*) induced at the specified concentrations of xylose and IPTG. **C. The increase in cell diameter caused by overexpression of *rodA* is reduced by simultaneous overexpression of *mreBCD*.** bMD557 (*yhdG*::Phyperspank-*rodA*::*cat amyE*::Pxyl-*mreBCD*::*erm)* was induced at the indicated levels of IPTG and xylose. **D. Rates of cell growth measured at the population (*Top*) and single cell (*Bottom*) level.** Mean rates of cell growth were measured either by *(Top*) optical density at 600 nm in a shaking plate reader in CH, or (*Bottom*) by microscopically assaying the rate single cells (grown under an agar CH pad) added surface area. All measures are from bMD620 grown in the inducer concentrations indicated along the bottom, with the exception of: “*ponA* null”, which is bMK005 (*ΔponA::cat*), “aPBP null”, which is bAM268 (Δ*pbpF*, Δ*pbpG*, Δ*pbpD*, Δ*ponA*::*kan*), and “aPBP Null + RodA”, which is bAM288 (Δ*pbpF* Δ*pbpG* Δ*pbpD* Δ*ponA*:*kan amyE*::Phyperspank*-rodA*-*His10*::*spec*), where RodA is induced with 0.05 mM IPTG. Dotted horizontal lines represent the mean growth rate of PY79 in CH medium, with the shaded area representing the SD.

Given that RodA and PBP2A must interact to synthesize PG ^30^, we tested if the widening caused by RodA overexpression required excess PBP2A. We created two strains that had *rodA* under xylose control; one contained native *pbpA*, and the other had *pbpA* under IPTG control (both as the only elongation-specific bPBP in the cell). As before, when *pbpA* was under native control, depletion or overexpression of *rodA* caused diameter to increase **(Figure 2B).** In contrast, when *pbpA* was held at the lowest induction required for WT width and *rodA* was simultaneously overexpressed, cells remained at WT widths. However, these RodA-overproducing cells could be made increasingly wider by increasing *pbpA* induction **(Figure 2B)**, demonstrating excess RodA requires excess PBP2A to increase width.

These results suggested RodA/PBP2A widens cells when their levels exceed the rest of the Rod complex. If this hypothesis is correct, wide cells caused by RodA/PBP2A overexpression should become thinner if MreBCD is increased, perhaps recruiting the diffusive excess RodA/PBP2A into circumferentially-moving, “thinning” Rod complexes. Indeed, while strong induction of *rodA* made wide cells, simultaneous overexpression of a second copy of *mreBCD* reduced cell diameter **(Figure 2C)**, a narrowing far greater than what we observed when overexpressing *mreBCD* in otherwise WT strains **(Figure 1C)**. Thus, the aPBP-like widening activity of RodA/PBP2A likely occurs once they exceed some level of MreBCD, possibly reflecting a saturation of binding sites within Rod complexes.

### Growth rates are maintained across a wide range of enzyme levels, unless both systems become limiting

We next examined how each PG synthetic system affected the rate of cell growth in our dual-induction strain using two assays: 1) single-cell measurements of the rate of surface area addition ^31^ under agar pads and 2) population measures of growth in a shaking plate reader. First, we examined the growth of the dual-inducible strain at the *mreBCD* and *ponA* inductions that produced WT widths **(Figure 1E)**. At the lowest induction pairs, growth was greatly reduced in both assays, and increased with each increasing induction pair **(Figure 2D)**, up to the induction pair that produced WT PBP1 and MreB protein levels. Thus, growth can be reduced if both PG synthesis systems become limiting.

Next, we assayed growth as we titrated either *mreBCD* or *ponA* while holding the other constant. Similar to *E. coli* MreB studies ^22, 32, 33^, both assays showed no difference in growth across a wide range of *mreBCD* inductions, except at the lowest induction where cells frequently lost rod shape. Similarly, both assays saw no difference in growth across our *ponA* induction range. Thus, even though these cells have different geometries, they add new surface area at the same rate. We next examined how extremely low levels of aPBPs affected growth. Similar to previous observations ^17, 34, 35^, both Δ*ponA* and ΔaPBP (Δ*pbpF*, Δ*pbpG*, Δ*pbpD*, Δ*ponA*::*kan*) strains showed a marked reduction in bulk growth, a defect rescued by overexpression of RodA ^6, 36^. However, the single cell measurements revealed a surprise: Both Δ*ponA* and ΔaPBP cells showed no defect in their single-cell growth rate, adding surface area at the same rate as WT, as did RodA overexpressing ΔaPBP cells **(Figure 2D)**. Given ΔaPBP cells display an increased rate of lysis that can be suppressed by RodA overexpression ^6, 36^, it appears the population growth rate defect of aPBP-deficient cells arises not from a reduction in growth rate, but from an increased frequency of death.

### Increased MreBCD increases the density of directionally moving MreB filaments and the fraction of directionally moving synthetic enzymes

To gain a mechanistic link between the level of the PG synthetic systems to cell width, we sought to develop microscopic measures of their activity. While the fraction of stationary aPBPs would be difficult to quantify, Rod complexes exhibit a more quantifiable phenotype: As their motion is driven by PG synthesis, the cellular activity of the Rod system can be measured by quantitating the number of directionally moving MreB filaments ^37^. We developed an analysis method that, using total internal reflection fluorescence microscopy (TIRFM) data, determines the density of MreB filaments moving directionally around the cell. Filaments undergoing directed motion are detected by taking advantage of the temporal correlation occurring between adjacent pixels across the cell width as objects move through them. These objects are then counted over a given time and then normalized to the total cellular surface area **(Figure 3A, S3A-F).** To evaluate the robustness of this approach to filament number, velocity, angle, and length, we simulated TIRFM time-lapses of MreB filaments in cells **(Figure S3G-K)**. This revealed this approach was more accurate than counting filaments in the same data with particle tracking. Furthermore, when applied to real TIRFM movies of MreB filaments, this approach yielded numbers comparable to tracking MreB filaments inside cells imaged with TIRF - structured illumination microscopy (TIRF-SIM) **(Figure S3L)**. With this analysis tool, we examined how the density of directionally moving MreB filaments related to rod width as we titrated the expression of an inducible *mreB-sw-msfGFP, mreCD* operon expressed as the only source at the native locus. This analysis revealed a well fit, apparently linear correlation between directional filament density and width (**Figure 3B**). Thus, the density of directionally moving MreB filaments reduces cell width, possibly by promoting directional PG synthesis.

**Figure 3.**
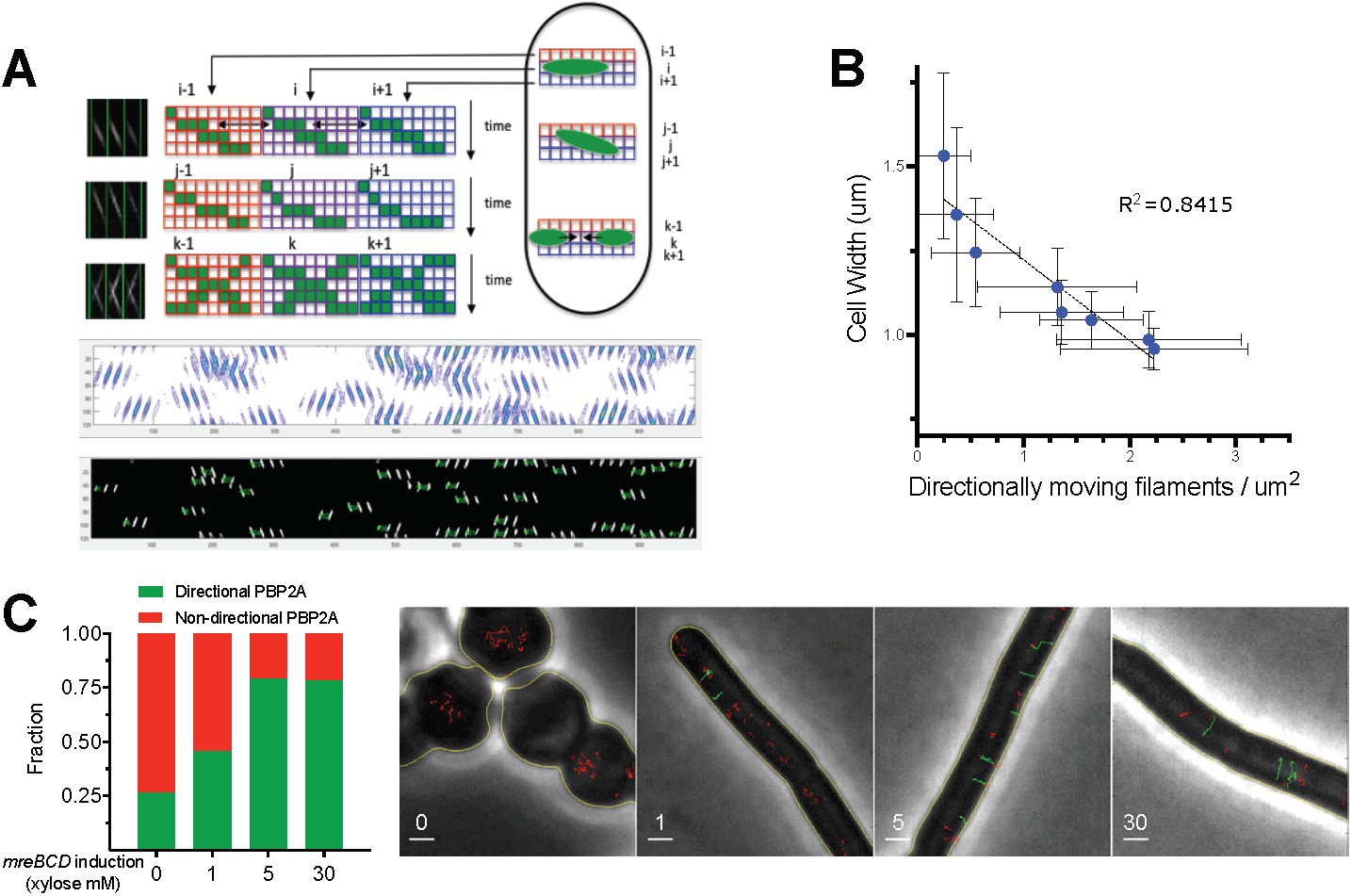
Increased *mreBCD* increases directional MreB filament density and the fraction of directional PBP2A molecules. **A. Schematic of the method to quantitate directionally moving MreB filaments.** Data shown is from simulated TIRFM time-lapses, with a 1 sec exposure time and 65 nm/pixel magnification. First, a kymograph is generated for each row of pixels along the midline of the cell. These kymographs are then lined up side by side to generate a single 2D image, where each column of the image contains a kymograph of each sequential row of pixels in the cell. This image is adaptively thresholded, then segmented with contour analysis to extract all fluorescent objects (middle). These objects are used to get velocity (slope), time (centroid), and position (row) for each particle. As each particle will show similar intensities in adjacent rows, or sometimes move at an angle, objects up to two rows apart are grouped based on time, position, and velocity. This yields the final particle count (bottom). See **Figure S3** for further details and tests. **B. The decreased rod width caused by increasing *mreBCD* induction correlates with an increasing density of directionally moving MreB filaments.** bYS981 (*amyE*::Pxyl*-mreB-SW-msfGFP, mreCD::erm ΔmreBCD::spec)* was grown in different amounts of xylose. Plotted are the steady state mean cell widths against the density of directionally moving filaments (determined as in **A**, above). All error bars are SD of the mean. **C. The fraction of directionally moving Halo-PBP2A molecules increases with *mreBCD* expression**. *mreBCD* was induced at the indicated levels in bMK385 (*amyE*::Pxyl-*mreBCD*::*erm* Δ*mreBCD::spec HaloTag-11aa-pbpA::cat*), and the single molecule motions of JF-549-labeled Halo-PBP2A were imaged by TIRFM using 300 msec continuous exposures. Plotted are the fractions of labeled PBP2A trajectories over 7 frames in length that moved directionally. *Right -* Representative montage of Halo-PBP2A trajectories at different levels of *mreBCD* inductions overlaid on phase images. Directionally moving tracks are labeled green; all other tracks are labeled red. Scale bar is 1 μm. See also corresponding **Movie S3**.

To this end, we next examined how MreBCD levels affected PBP2A motion. Multiple studies have noted PBP2A molecules exist as a mixed population; some diffuse within the membrane, while others move directionally ^12^-^14^. We titrated *mreBCD* induction as we tracked the motions of PBP2A molecules (expressed at the native locus as a HaloTag fusion, sparsely labeled with JF-549 ^38^). At low *mreBCD*, only a small fraction of PBP2A moved directionally. As we increased *mreBCD* induction, an increasing fraction of PBP2A molecules moved directionally **(Figure 3C, Movie SM1)**, demonstrating that *mreBCD* limits the amount of directional PBP2A.

### Increased MreBCD:PBP1 correlates with an increased amount of oriented material and structural anisotropy of the cell wall

To gain more insight into how an increased density of directional rod complexes can reduce rod width, we used polarization microscopy to understand how increased circumferential synthesis affected the organization of material within the cell wall. Polarization microscopy reports on both the angle and extent of orientation within optically anisotropic (or birefringent) materials ^39^, and has been used to assay the orientation of various materials, including plant cell walls ^40^-^42^ and mitotic spindles ^43^. Polarization microscopy revealed purified WT sacculi show birefringence, indicating that some fraction of the material within them is oriented. Focused at their surface, sacculi showed a predominant slow axis oriented along the rod length (**Figure 4A, Movie SM2**). Given that amino acids have a higher refractive index compared to sugars, this suggests (in agreement with previous models ^44^) the peptide crosslinks are predominantly oriented along the rod length, and the glycans are oriented around the circumference.

**Figure 4.**
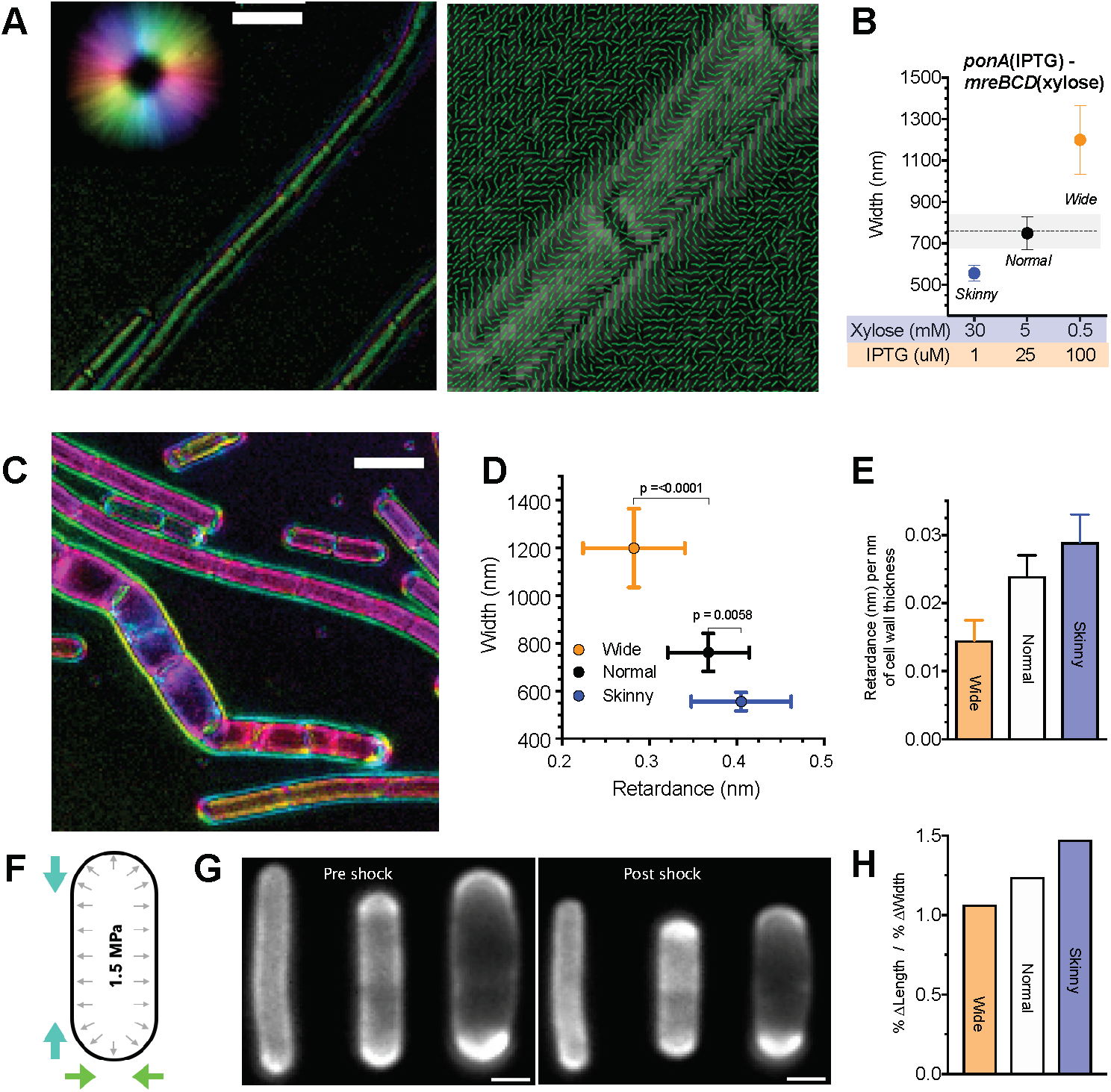
Increased Rod activity increases both the amount of oriented material within sacculi and their mechanical anisotropy. All error bars are SD of the mean. **A. Polarization microscopy reveals oriented material within the cell wall.** *Left* - LC-PolScope images of purified WT PY79 sacculi. Focused at the surface, the cell wall is seen to be birefringent. Color indicates the orientation of the slow axis, with intensity corresponding to retardance in that direction. Orientation key is the color wheel at upper left. *Right* - Polarization orientation view of the sacculi surface, with lines pointing in the predominant orientation of the slow axis. See also **Movie S2.** **B. Induction conditions used to assay sacculi properties.** bMD620 (*amyE*::Pxyl*-mreBCD*::*erm* Δ*mreBCD*::*spec yhdG*::Pspank*-ponA*::*cat* Δ*ponA*::*kan*) was induced to grow at 3 different widths: Skinny (30 mM xylose, 1 μM IPTG), Normal (5 mM xylose, 25 μM IPTG), and Wide (0.5 mM xylose, 100 μM IPTG). **C.** Example LC-PolScope image of bMD620 sacculi induced at different widths. For each experiment, pairs of purified sacculi (“Wide” and “Normal”, or “Wide” and “Skinny”) were combined, and Z-stacks were collected in 100 nm steps. See also **Movie S3.** **D. The amount of oriented material in the cell wall increases with *mreBCD* induction, and inversely correlates with rod width.** Plotted is the mean retardance vs. width along the length of projected Z-stacks of at least 90 different cells for each induction condition. **E.** Mean retardance for each sample normalized to the thickness of the cell wall, as measured by TEM. **F. Schematic of osmotic shock assay of anisotropy.** *B. subtilis* sacculi are normally under tension due to the high internal turgor (black arrows). Hyperosmotic shock negates this pressure, allowing observation of how sacculi shrink in length and width (colored arrows). **G.** Example FDAA-labeled cells from each condition (*left to right*: Skinny, Normal, Wide) before and after osmotic shock. Scale bar is 1 μm. **H. As the relative amount of Rod system activity increases, so does the mechanical anisotropy of the sacculus.** Anisotropy (% change in length / % change in width) for each induction condition following osmotic shock. Raw data and reduction for each dimension (both absolute and percent) are in **Figure S4**.

We then examined how the balance between the Rod and aPBP systems affected the relative amount of oriented material in the wall. We grew our dual-induction strain under three different induction conditions to create varied width cells: wide (high *ponA*, low *mreBCD*), normal (*ponA* and *mreBCD* induced at WT levels), and skinny (low *ponA*, high *mreBCD*) (**Figure 4B**), purified their sacculi, and quantified their total retardance with polarization microscopy (**Figure 4C, Movie SM3**). Retardance is the differential optical path length for light polarized parallel and perpendicular to the axis of molecular alignment; alternatively, it is defined as birefringence (Δ*n*) times the physical path length through an anisotropic material. Wide cells (high *ponA*, low *mreBCD*) had the lowest retardance, skinny cells (low *ponA*, high *mreBCD*) had the highest retardance, and normal cells were in between (**Figure 4D**). Because retardance depends on both the thickness of material (path length) and the degree of its orientation (Δ*n*), we normalized the retardance of each sample to the mean thickness of the cell walls of each induction condition, obtained using transmission electron microscopy (**Figure S4A)**. This revealed that the skinny cell walls had more highly ordered material (Δ*n*) per path length (nm) of cell wall thickness, and that wide cell walls had the least (**Figure 4E**). Thus, in agreement with recent atomic force microscopy studies showing orientated glycans in *E. coli* require MreB ^16^, these experiments demonstrate that as Rod system activity increases, so does the amount of oriented material in the wall.

The sacculi of *E. coli and B. subtilis* are mechanically anisotropic, stretching more along their length than across their width ^44^-^46^. To test how the ratio of oriented to unoriented PG synthesis affected this property, we grew our dual-inducer strain at the three *mreBCD:ponA* inductions above, labeled their walls with Alexa-488-D-amino carboxamides ^4^, and assayed the dimensions of their cell walls before and after hyperosmotic shocks (**Figure 4F-G**). This revealed that increased Rod system activity correlated with an increased mechanical anisotropy of the sacculus: As we increased the expression of MreBCD relative to PBP1A, rods shrank less across their width, and more along their length **(Figure 4H, S4B-C)**. Thus, the Rod system acts to reinforce rod shape against internal turgor by promoting oriented PG synthesis around the rod.

### *E. coli* Rod mutants also show a correlation between cell width and the density of directionally moving filaments

Previous studies have examined how MreB and PBP2 mutations affect the shape of *E. coli*, hypothesizing that their abnormal widths arise from changes in the curvature, twist, or angle of MreB filaments. Our observations in *B. subtilis* suggested an alternative explanation: Abnormal width might arise from simply changing the amount of Rod system activity. We tested this by measuring the density of directional GFP-MreB filaments in these same mutant *E. coli* strains. As a benchmark, we first assayed the width of *E.coli* as we titrated the expression of mreB-sw-msfGFP using CRISPRi against msfGFP ^47^ ^28^. As in *B. subtilis*, this yielded an inverse relationship between directional MreB filament density and cell width (**Figure 5A)**. Examining each group of mutants showed the same result: 1) An identical trend was observed for the mutations hypothesized to change filament twist ^19^, 2) as well as in the mutations designed to change filament curvature ^24^. And 3) notably, the same correlation between directional filament density and cell width was observed in strains where *E. coli mrdA* (PBP2) was replaced with *mrdA* genes from other species ^22^. TIRF-SIM imaging of these MreB mutants revealed some insight into these effects (**Movie SM4**): Some of the wide mutants showed either A) longer but fewer filaments, or B) a large fraction of immobile filaments. Conversely, some thinner mutants appeared to have more, but shorter filaments. Finally 4), while Colavin et al. hypothesized that RodZ reduces width by changing MreB filament curvature ^24^, we found that increased RodZ induction increased MreB filament density as cells thinned (**Figure 5B, Movie SM5**). This suggests that the decrease in cell width most likely occurs from to RodZ’s ability to nucleate MreB filaments ^48^ rather than changing their curvature. Thus, the density of directionally moving Rod complexes correlates with cell width in both *B. subtilis* and *E. coli* across multiple genetic perturbations.

**Figure 5.**
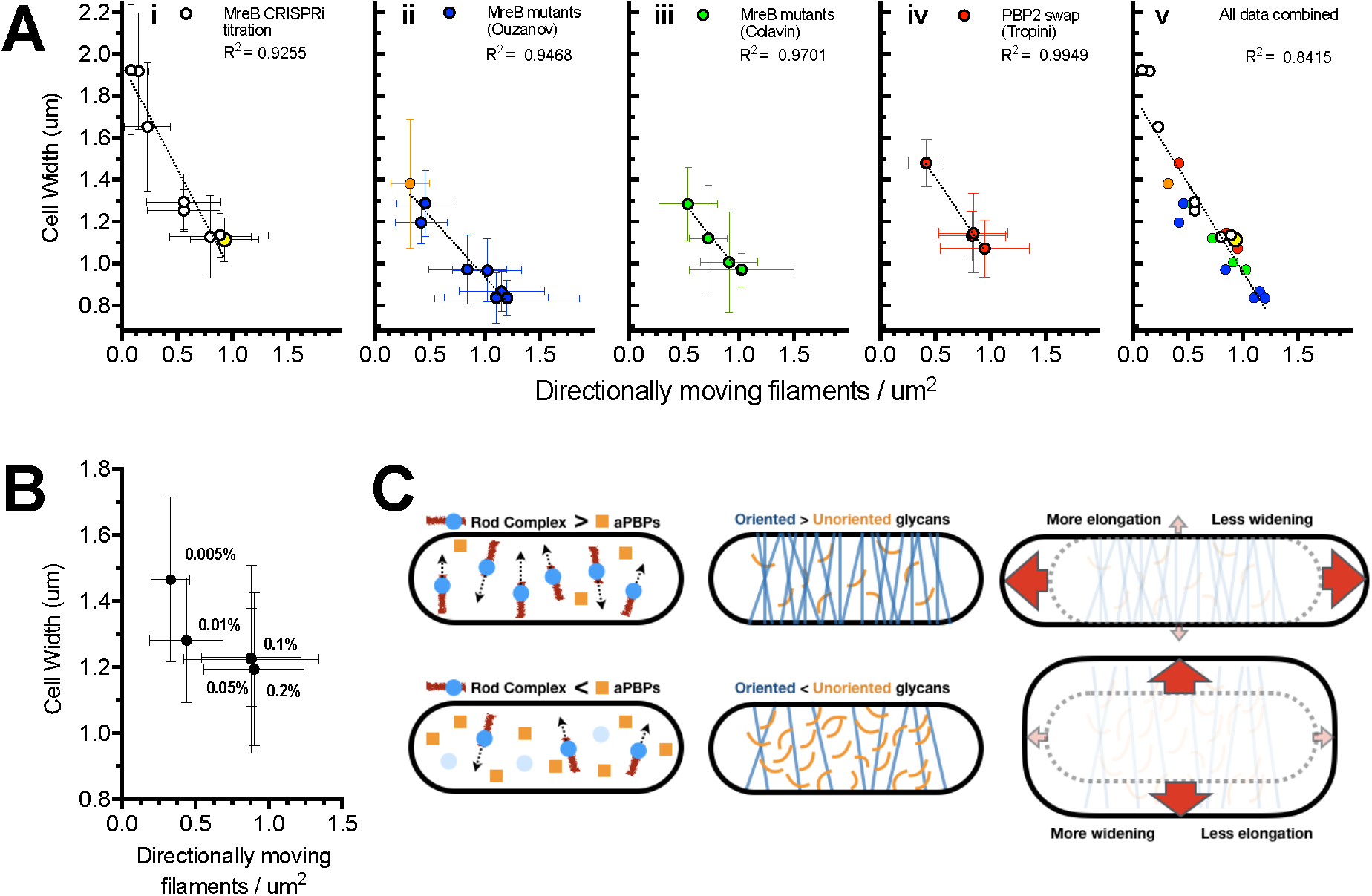
– Directional MreB filament density also correlates with cell width of *E. coli* Rod mutants, suggesting a generalizable model for the effects of Rod complex on cell width. **A. Cell width vs. density of directionally moving MreB filaments in different *E. coli* strains. (i)** AV88 (186::P_tet_-*dCas9*, *mreB*::*msfGFP*-*mreBsw*), allows the tunable expression of *mreB-msfGFPsw* by expressing various sgRNAs with different matches against msfGFP. Yellow indicates WT expression. **(ii)** mreB-SW-msfGFP mutant strains from Ouzounov et al., 2016. Orange indicates RM478 *(ΔrodZ, mreB(S14A)-msfGFPsw)* from Morgenstein et al., 2015. **(iii)** mreB-msfGFPsw mutants believed to change filament curvature from Colavin et al., 2018. **(iv)** msfGFP-mreB strains from Tropini et al., 2015, where *mrdA* is replaced with *mrdA* from other species. **(v)** All data from (i)-(iv) combined. See also **Movie SM4**. All values are detailed in **Table S1**. **B. Decreased cell width caused by increased RodZ expression correlates with an increased density of directionally moving MreB filaments.** KC717 (*csrD::kan, mreB::msfGFP-mreB, ProdZ<>*(*frt araC PBAD*)) was grown at different arabinose concentrations (indicated on the graph), and filament density was calculated as in **Figure 3B**. See also **Movie SM5**. **C. Model for how the balance between aPBPs and the Rod system affects cell width.** *Left* – When Rod complex activity (perhaps set by the number of MreB filaments) is high relative to that of aPBPs, sacculi have more circumferentially oriented material (*Center*) compared to when aPBP activity is greater. *Right* – As the amount of oriented material increases, the less elastic sacculi are more rigid across their width, but less rigid along their length. Being stretched by the internal turgor pressure, sacculi with greater Rod activity are better able to maintain their width, while stretching more along their length. In contrast, cells with reduced Rod activity have less circumferentially oriented glycans to reinforce their width, and thus expand more along their width.

## Discussion

The shape of bacteria is defined by their cell walls; these experiments demonstrate that the two systems that insert PG into it have opposing roles on its shape. Due to the intrinsic orienting of MreB filaments around the rod width ^11^, the Rod system inserts circumferentially oriented material around the rod circumference, reducing its diameter. As the number of MreB filaments increases, so does the fraction of directional enzymes and the amount of oriented material in the wall. In contrast, the aPBPs do not move circumferentially, inserting material that isotropically enlarges the sacculus. Our data indicates the macroscopic shape of the sacculus arises from the nature of the material inserted into it: The more it is oriented around the rod circumference by the Rod complex, the less the rod stretches across its width, and the more it stretches along its length **(Figure 5C)**.

If the balance between the two PG synthetic systems is perturbed, the shape of the sacculus becomes altered, though its rate of expansion remains constant. As both systems utilize the same pool of lipid II ^49^, the flux through each may depend on their relative levels; if one is reduced, the flux through the other may increase. This would explain why disrupting the Rod system causes cells to swell ^27, 29, 50^-^52^: In the absence of Rod-mediated thinning, aPBPs add more material uniformly over the cell surface. Likewise, in the absence of aPBP-mediated widening, increased flux through the Rod system would explain why cells become extremely thin ^17, 53^. However, if both systems are equivalently reduced, cells grow with normal widths, but at slower rates; as long as the activities are balanced - identical shape arises from the balanced levels of enzymes, but growth is reduced due to their combined activity becoming limiting. This would explain why *ponA* mutations rescue *mreB* deletions ^54^; equally crippling both systems may rebalance the activities such that the cell retains normal shape and viability.

### Implications for the role of MreB in rod width determination

Given that 1) *mreBCD* from the wide bacterium *B. megaterium* creates close to normal diameter *B. subtilis* rods, and 2) *B. subtilis* diameter depends on *mreBCD* levels, we find it unlikely that any property of MreB filaments defines a given cell diameter. Rather, MreB appears to be one component of a rod-thinning system, working in opposition to aPBP-mediated widening. Indeed*, in vitro* studies have revealed MreB filaments are extremely curved (> 200 nm), allowing them to deform to, and orient around the width of bacteria of any larger width ^9, 11^. While it remains possible that given MreB mutations or interactions with RodZ could indeed alter filament curvature or twist ^19, 21, 24, 26^, these experiments suggest a more parsimonious explanation for their effect on Rod activity: These mutations alter MreB’s polymerization dynamics or cellular distribution. Reducing the amount or number of active MreB polymers would reduce Rod activity, causing cells to widen. Likewise, mutations altering MreB’s polymer length distribution or tendency for filaments to bundle would cause the distribution of Rod activity to become non-uniform, causing some parts of the cell to thin while others would widen, as previously observed for certain MreB mutants ^12, 27^.

Additionally, our experiments reinforce observations that aPBPs and RodA (when in excess to MreBCD) serve reparative, anti-lytic roles: aPBP-deficient cells grow at the same rate as WT, yet have an increased frequency of death, lysing as thin rods without losing shape ^6, 36^. Likewise, aPBP-mediated synthesis increases upon endopeptidase overexpression in *E. coli*^55^. Given the active state of aPBPs correlates with single aPBP molecules displaying periods of transient immobility ^14, 56^, their synthesis may be localized to small (<50 nm) regions. Combined, these observations support a model where aPBPs synthesize material to fill gaps in the PG meshwork ^36, 14^. Gaps could arise via mechanical damage, hydrolases ^55^, or between the imperfectly oriented strands built by the Rod complex ^11^. If this model is correct, the different spatial activities of these two systems might allow the sacculus to maintain integrity at any Rod/aPBP ratio: the fewer the Rod complexes, the larger the gaps filled by aPBPs.

While many different activities affect the shape of the sacculus, such as its cleavage by hydrolases or rigidification by wall teichoic acids ^11^, the first process defining its geometry is the spatial coordination controlling where glycans are inserted into it. While these experiments give a coarse-grained description into how each synthetic system affects cell shape, a fine scale, mechanistic understanding remains to be determined - How does aPBP activity make cells wider? How does increased Rod activity make cells thinner? Understanding the physical mechanisms causing these changes will require not only a better understanding of the molecular architecture of the sacculus, but also an investigation into how the enzymes downstream of glycan insertion affect the shape and mechanics of sacculi as they subsequently modify, remodel, and break down the nascently-polymerized glycan architecture.

## Acknowledgements

We would like to thank Carl Wivagg, Ye Jin Eun, Leigh Harris, and Thomas Bernhardt for helpful advice and discussions, Georgia Squyres for reading of the manuscript and TIRF-SIM acquisition, and Luke Lavis for his generous gift of JF dyes. We thank Z. Gitai and K.C. Huang for providing strains. TIRF-SIM was performed at the Advanced Imaging Center at Janelia Research Campus, a facility jointly supported by the Gordon and Betty Moore Foundation and Howard Hughes Medical Institute. This work was funded by National Institutes of Health Grants R01GM114274 to RO, and DP2AI117923-01 to EG, as well as a Smith Family Award, Searle Scholar Fellowship, and support from the Volkswagen Foundation to EG and SVT. This work was performed in part at the Center for Nanoscale Systems at Harvard University, supported by NSF ECS-0335765.

## Author Contributions

*B. subtilis* strains were cloned by MD, YS, and MK. All width and bulk growth measurements of *B. subtilis* were done by MD, and *E. coli* widths by YS. *E. coli* CRISPRi strains were cloned by AV, who was supervised by SVT. Single cell growth rates were done by YS and MK. TIRFM and tracking of PBP2A was done by MK. All TIRFM of MreB was done by YS. TIRF-SIM of MreB was done by YS, EG, and JR. Purified sacculi and proteomic mass spec sample preps were done by MD. YS wrote the code for analysis of single cell growth rate, filament density, and simulations of data. Polarization microscopy and analysis was conducted by JR, RO, and EG. SW did the FDAA synthesis, osmotic shocks, and TEM. The paper was written by EG, MK, MD, and YS.

## Supplemental Material for

### Supplemental figures

**Figure S1.**
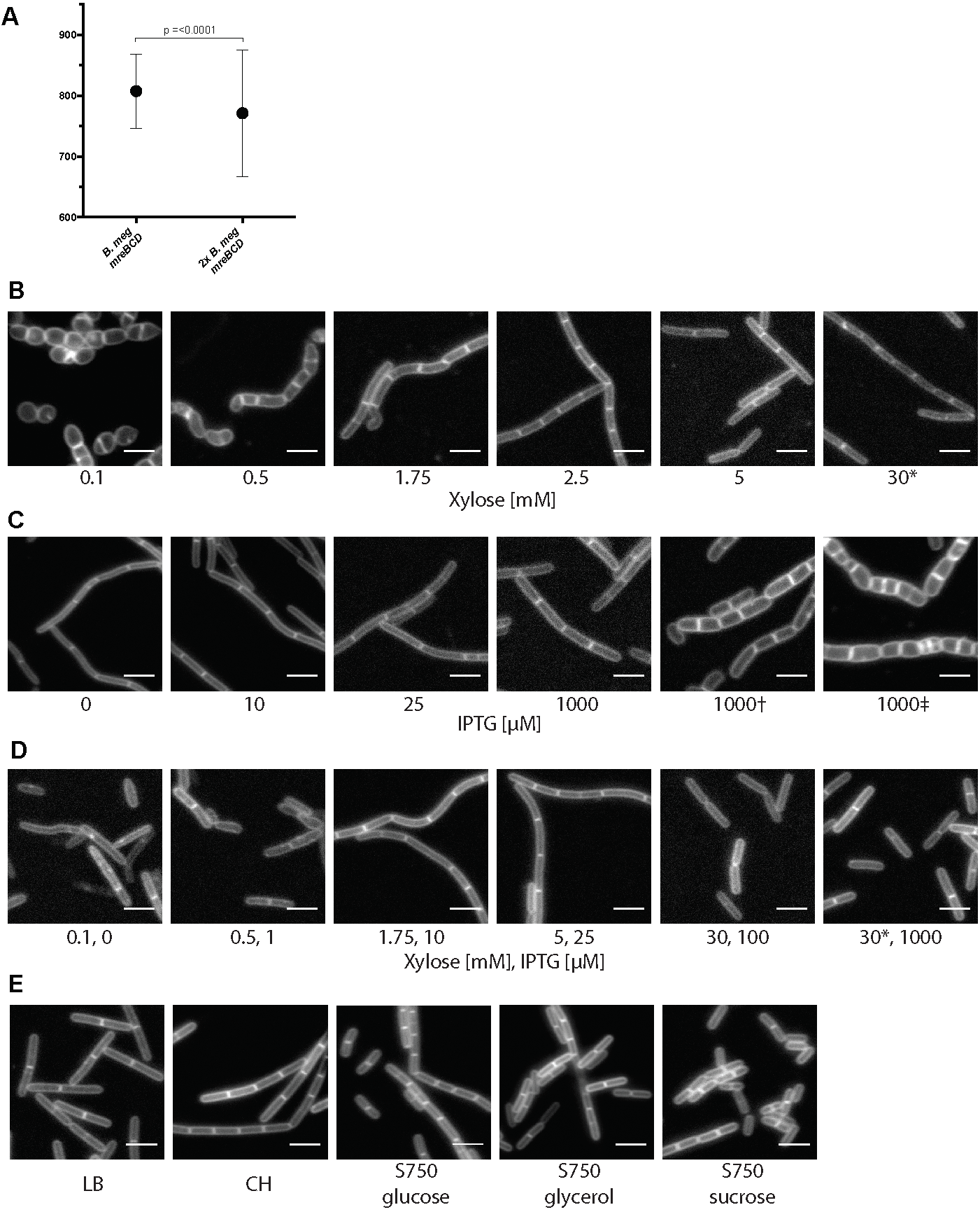
**A.** Zoomed view of the two rightmost distributions of widths in **Figure 1B**. Error bars are SD of the mean. **B-D.** Cells were grown in CH media with the indicated concentrations of inducers during steady state growth, then stained with 0.5 μg/mL FM 5-95. Scale bars are 4 μm. **B.** Images are of bMD545 (*amyE::Pxyl-mreBCD::erm, ΔmreBCD::spec)*, save the induction annotated * which is bMK355 (*amyE::Pxyl-mreBCD::erm*), containing a xylose inducible *mreBCD* in addition to the native *mreBCD*. **C.** Images are of bMD598 (*yhdG::Pspank-ponA::cat, ΔponA::kan*), induced with the indicated amounts of IPTG, save the inductions marked † and ‡ which are under stronger promoters than bMD598; † is bMD586 (*yhdG*::Phyperspank-*ponA*::*cat*, Δ*ponA::kan*), and ‡ is bMD554 (*yhdG*::Phyperspank-*ponA*::*cat*), containing an inducible *ponA* in addition to the native copy. **D.** Images of bMD620 (*amyE*::Pxyl-*mreBCD*::*erm*, Δ*mreBCD::spec, yhdH*::Pspank-*ponA*::*cat*, Δ*ponA::kan*) grown in CH, and induced with the indicated amounts of IPTG and xylose. The image annotated **\*** indicates bMD622 (*amyE::Pxyl-mreBCD*::*erm yhdH::Pspank*-*ponA*::*cat* Δ*ponA::kan*) containing a xylose-inducible *mreBCD* in addition to the native *mreBCD.* **E.** Representative images of PY79 grown in indicated media, then stained with 0.5 μg/mL FM 5-95. Scale bars are 4 μm.

**Figure S3.**
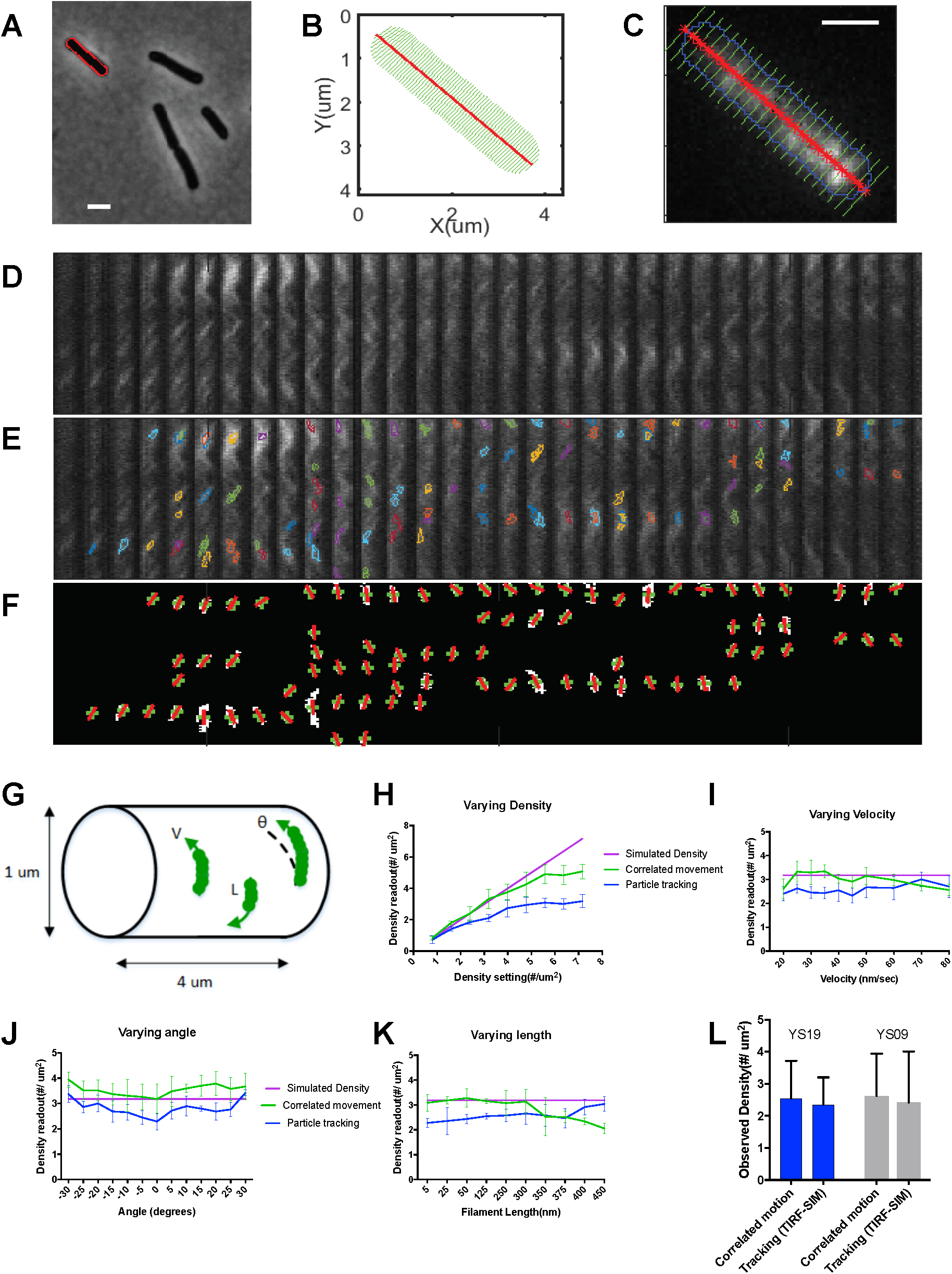
A-F. Expanded description of the method used to determine directional filament density. Cells expressing fluorescent MreB fusions were imaged with TIRF for 120 seconds, followed by a single image in phase-contrast. From the phase-contrast images (**(A) -** scale bar is 2 um), cells were segmented to determine their dimensions and midline **(B)**. The resulting segmentation mask was used to create kymographs out of the fluorescence time-lapse data. **(C)** Shows the mask overlaid on a maximum intensity projection of a TIRF time-lapse (red line indicates midline, green lines are the axis along which kymographs are drawn, scale bar is 2 um). Next, kymographs were created for each sequential row of pixels along the midline, and the kymographs from each row were displayed side by side as in **(D)**. Next, objects within these kymographs were identified and segmented based on contour analysis **(E)**, using adaptive thresholding. For each identified object, a set of properties [*X, t, v, I*] were measured. (*X, t*) are the centroid of the object in the binary image (green crosses in **(F)**). *X* is the position of the filament along the cell width, and *t* is time, corresponding to rows in the kymographs. Velocity (*v*) is calculated based on the diagonal slope of the object in the binary image (red lines in **(E**)), and can be positive or negative depending on the direction of motion. Intensity (*I*) is the sum of the intensities of all pixels in the object. Because each kymograph represents one row of pixels in the cell, and the signal from each particle can span more than one pixel (65 nm) in width, the same filament can appear in several sequential kymographs (or rows in the original image). In order to link the objects occurring in multiple rows together, each object in a given row [*X*_i_] can be linked to corresponding objects in the next row [*X*_i+1_] or previous row [*X*_i-1_] based on their time and velocity. Objects are only linked if they have a velocity *v* +/-20 nm/sec of each other, and their peak time *t* is within 2 seconds; these objects are then counted as a single filament. This also allows us to capture filaments moving at angles of up to +/- 30 degrees to the midline. Finally, the number of counts occurring during the imaging interval is normalized based on the cell width and length (or total surface area Π*width*length). **G-J. Evaluation of filament density quantitation tested using simulations.** **(G)** Schematic of the MreB data simulations. Simulations were run using custom MATLAB code, with the following parameters: Maximum fluorescence intensity was 300 counts and noise was 10 (approximating the signal to noise conditions in our imaging). The simulated cell had a width of 1 μm and a length of 4 μm. We tested the robustness of our tracking by varying one parameter while the others were fixed at the following values: Filament density was 3.2/μm^2^ (corresponding to 40 MreB filaments per cell), filament velocity was 30 nm/sec, filament orientation was 0 degrees, and filament length was 250 nm. This data was tracked using the correlated motion method detailed above. For comparison, we tracked the same simulated data with the linear motion LAP tracker in Trackmate, with a low bar for calling a trace a directional motion (tracks longer than 6 frames with a displacement greater than 250 nm). **(H)** Filament detection performance with respect to different filament density settings (from 0.8 to 7.2/um^2^, corresponding to 10 - 90 MreB filaments per cell). **(I)** Filament detection with respect to filament angles (from −30 to 30 degrees). **(J)** Filament detection with respect to different filament velocities (from 20 to 90 nm/sec). **(K)** Filament detection with respect to different filament lengths (5 nm to 450 nm). **L. Correlated motion analysis yields a similar density of directional filaments as tracking of MreB filaments imaged by TIRF-SIM.** For this comparison, two different strains with natively expressed MreB fusions, YS09 (*mreB-mNeonGreen*) and YS19 (*mreB-SW-msfGFP*), were grown under the same conditions, and were analysed using two different imaging/analysis pipelines: 1) As above, standard TIRFM imaging followed by correlated motion analysis, and 2) TIRF-SIM imaging, followed by tracking filaments with particle tracking (using the same settings as in **(G)** above).

**Figure S4.**
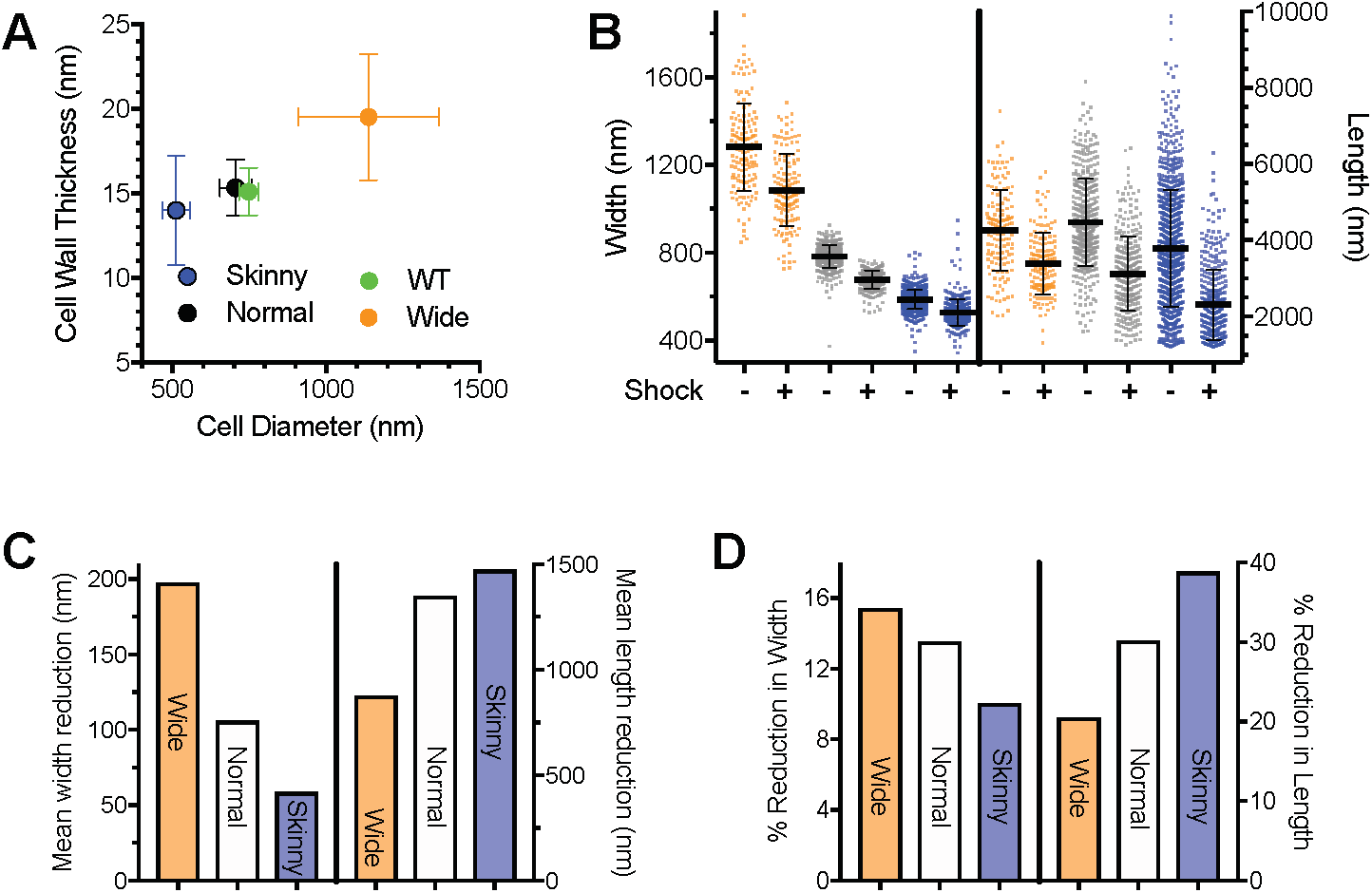
**A.** Cells, induced under identical conditions to **Figure 4B**, were fixed and stained with uranyl acetate, the block was cut in ultrathin sections, then imaged with transmission electron microscopy (TEM). The thickness of the cell wall was measured and plotted against cell diameter (also determined via TEM). **B.** All data of the width (left) and length (right) of cells before and after osmotic shocks (orange = Wide, grey = Normal, blue = Skinny). Error bars are SD of the mean. **C** Absolute and **D** percent reduction in width and length of labeled sacculi under each induction condition after osmotic shock.

## Supplemental Movie Legends

### Movie S1

Example single molecule movies of PBP2A-HaloTag (labeled with JF549^1^) at different *mreBCD* induction levels in bMK385 (*amyE::Pxyl-mreBCD*::*erm, ΔmreBCD, HaloTag-11aa-pbpA::cat*). Xylose concentrations are indicated in each panel. Frames are 300 msec apart.

### Movie S2

Z-stack of LC-PolScope images of sacculi purified from WT strain PY79. (Left) Z-stack of the retardance of each plane. (Right) Color map of birefringence at each plane, with color wheel at upper left giving orientation reference. Z-steps were taken in 100 nm increments.

### Movie S3

Z-stack color map of birefringence of sacculi purified from strain bMD620 (*amyE::Pxyl-mreBCD::erm, ΔmreBCD, yhdG::Pspank-ponA::cat*, *ΔponA::kan)*, where cells were induced to “Wide” (0.5 mM xylose, 0.1 mM IPTG) and “Normal” widths (5 mM xylose, 0.025 mM IPTG). Color wheel at upper left indicates orientation direction. Z-steps were taken in 100 nm increments.

### Movie S4

TIRF-SIM movies of different mreB-SW-msfGFP mutants. Cells were grown in LB at 37°C, then placed under an LB agar and imaged at 37°C. Frames are 1 sec apart. Scale bars are 1 um.

### Movie S5

TIRF-SIM movies of strain KC717(*mreB::msfGFP-mreB, ProdZ<>*(*frt araC PBAD*)). Cells were grown for 5 hours in LB with the indicated arabinose concentrations, were placed under an LB agar pad and imaged at 37°C. Frames are 1 sec apart. Scale bars are 1 um.

## Materials and Methods

### Media and culture conditions

For all experiments, unless otherwise noted, *B. subtilis* and *B. megaterium* were grown in casein hydrolysate (CH) medium (where indicated, xylose and/or isopropyl thiogalactoside (IPTG) was added), and *E. coli* strains were grown in lysogeny broth (LB) medium (where indicated, arabinose or anhydrotetracycline (ATc) was added), at 37°C with rotation. When pre-cultures, grown overnight at 25°C from single colonies, reached exponential phase the next day, they were further diluted into fresh growth medium (and where indicated, with the specified concentrations of inducer) and were grown for at least 3 hours at 37°C with rotation, to an OD_600_ of ∼0.3 to 0.7.

### Sacculi purification

20 mL cultures were grown in 250 mL baffled flasks at 37°C with vigorous shaking, to an OD_600_ of ∼0.5, were harvested and were centrifuged at 5,000 × *g* for 5 min at 4°C. Cell pellets were resuspended in 1 mL of ice-cold phosphate buffered saline (PBS), were centrifuged at 6,000 × *g* for 30 sec, were resuspended in 500 μL of PBS, and were killed by boiling in a water bath for 10 min. Cells were centrifuged at 6,000 × *g* for 2 min, were resuspended in 500 μL of PBS containing 5% sodium dodecyl sulfate (SDS), and were boiled in a water bath for 25 min; this was repeated once, except boiling was for 15 min. To remove the SDS, the samples were centrifuged at 6,000 × *g* for 2 min, and were resuspended in 500 μL of PBS; this was repeated 5 times. The cells were centrifuged at 6,000 × *g* for 2 min, were resuspended in 1 mL of 50 mM Tris-HCl, pH 7.5, containing 10 mM NaCl, and 2 mg of pronase from *Streptomyces griseus* (MilliporeSigma, MA), and were incubated at 60°C for 90 min with gentle shaking. To remove the pronase, the samples were boiled twice in PBS/5% SDS, followed by 6 rounds of washes in PBS, exactly as described in the steps above that precede the pronase treatment. The samples were centrifuged at 6,000 × *g* for 2 min, were resuspended in 1 mL of 25 mM Tris-HCl, pH 8.5, containing 0.5 M NaCl, 5 mM MgCl_2_, and 100 U of salt-active nuclease (SAN) (ArticZymes, Norway), and were incubated at 4°C overnight with gentle mixing. To remove the SAN, the samples were boiled twice in PBS/5% SDS, followed by 6 rounds of washes in PBS, exactly as described in the steps above that precede the pronase treatment. The sacculi were centrifuged at 6,000 × *g* for 2 min, the supernatant was removed, and the pellets were stored at −80°C.

### Polarization microscopy

Purified sacculi were resuspended in PBS and placed on ethanol-cleaned No. 1.5 coverslips under a cleaned glass slide. Polarization images were acquired on an inverted LC-PolScope mounted on a Nikon Ti-E equipped with a 60x/1.4NA Plan Apo oil immersion objective and oil condenser with matching NA, and a Hamamatsu Photonics Flash4 camera. Z-stacks in 100 nm steps were taken of each sample using 50 msec acquisitions of 546/12 nm light. All image acquisition, processing, and display, including colored display and line maps were prepared using the OpenPolScope Hardware Kit and plugins for ImageJ/Micro-Manager from OpenPolScope.org. From the previously prepared sacculi, multiple slides were imaged and quantitated, with independent background calibrations. These gave similar results and were combined to yield the final data set.

### Calculation of retardance

Z-stacks of the computed total retardance for each slice were exported from OpenPolScope. These stacks were cropped at one frame above and beneath the focal planes of the top and the bottom of the cells, then projected into a single plane using ImageJ. To avoid getting high retardance values from the edge effects arising from the sides of the cell or the septa, we selected the retardance at the middle of the cell, using the average of line scans (5 pixels long and 3 wide) that were drawn down the center of cells, taking care to avoid edges and septa.

### Transmission electron microscopy of cell wall thickness

Overnight cultures were diluted into fresh medium and grown to an OD_600_ of ∼0.3 to 0.5. Cells were pelleted by centrifugation at 5,000 x *g* and fixed by resuspending in 100 mM MOPS pH 7 containing 2% paraformaldayde, 2.5% gluteraldehyde, and 1% DMSO overnight at 4°C. Cells were centrifuged at 5,000 x *g* and were washed 3 times with 100 mM MOPS pH 7. The pellet was stained with 2% osmium tetroxide in 100 mM MOPS for 1 hr, washed twice with deionized water, and stained overnight with 2% uranyl acetate. The pellet was washed twice with deionized water and dehydrated by washing once with 50% ethanol, once with 70% ethanol, once with 95% ethanol, and then three times with 100% (v/v) ethanol. Samples were prepared for resin infiltration by washing once with 50% ethanol, 50% propylene oxide (v/v) and then once with 100% propylene oxide. All wash steps were for 5 min. Infiltration of resin was achieved by incubation with 50% Embed 812 (EMS, PA)/50% propylene oxide for 1 hr, followed by incubation with 67% Embed 812/33% propylene oxide for 1hr, and incubation with 80% Embed 812/20% propylene oxide for 1 hr. Samples were then incubated twice with Embed 812 for 1 hr, followed by an overnight incubation in molds. The molds were baked at 65°C for 18 hr before sectioning.

Serial ultrathin sections (80 nm) were cut with a Diatome diamond knife (EMS, PA) on a Leica Ultracut UCT (Leica Microsystems, Germany) and collected on 200-mesh thin-bar formvar carbon grids. Sections were imaged on an FEI Tecnai transmission electron microscope.

Cell wall thickness measurements were performed using a custom-built MATLAB (Mathworks, MA) script. Image intensity profiles extracted from lines were drawn perpendicular to a user-input line defining the middle of the cell wall. The distance between the two lowest points below a threshold within 40 nm of the middle of the cell wall was measured as the cell wall thickness at ∼30 points in each cell. This experiment was conducted once, using multiple cells for the analysis.

### Measurements of *Bacillus* cell dimensions

Cultures grown to an OD_600_ of ∼0.3 to 0.7 were stained with 0.5 μg/mL FM 5-95 (Thermo Fisher, MA) for 1 min, and were concentrated by centrifugation at 6,000 x *g* for 30 sec. The cell pellet was resuspended in ∼1/20 volume of growth medium, and 3 μL was applied to ethanol-cleaned No. 1.5 coverslips under a 3% agarose pad containing growth medium. Fluorescent cells were imaged with the top surface of the agarose pad exposed to air, in a chamber heated to 37°C. Epifluorescence microscopy was performed using a Nikon Eclipse Ti equipped with a Nikon Plan Apo λ 100×/1.4NA objective and an Andor camera. Cell contours and dimensions were calculated using the Morphometrics software package ^2^. Each “Width” data point (**Figures 1-4)** is calculated from at least 79 cells, but most typically hundreds (see **Table S1**) from multiple fields of view across different areas of the agarose pad. Key points in these experiments were repeated multiple times on independent days; including induction conditions for bMD545, bMD598, and bMD620 that resulted in WT width and the extremes of thinning and widening; and PY79 measured in parallel as a width control. All repeat measurements gave similar mean values.

### Measurements of single-cell growth rate

Cultures grown to an OD_600_ below 0.3 were concentrated by centrifugation at 6,000 x *g* for 30 sec. The cell pellet was resuspended in growth medium, and applied to No. 1.5 glass-bottomed dishes (MatTek Corp., MA). All cells were imaged under a 2% agarose pad containing growth medium, with the top surface exposed to air, in a chamber heated to 37°C. Phase-contrast microscopy was performed using a Nikon Eclipse Ti equipped with a Nikon Plan Apo λ 100×/1.4NA objective and an Andor camera. We used a custom-built package in MATLAB to perform segmentation on phase-contrast time-lapse movies, then calculated the growth rate of the surface area of single *B. subtilis* chains. Each data point for the single-cell growth rates (**Figure 2D**) is the result of a single experiment; for each, >50 cells from multiple fields of view across different areas of the agarose pad were imaged and analyzed.

### Measurements of bulk growth rate

For cell culture measurements of growth rate, overnight pre-cultures in mid-log growth were diluted in fresh medium and grown for ∼3 hr at 37°C to an OD_600_ of ∼0.3 to 0.7. The cultures were diluted back to a calculated OD_600_ of 0.07 in 100-well microtiter plates (replicated in 3 to 5 wells for each culture), and their growth rates were measured in a Bioscreen-C Automated Growth Curve Analysis System (Growth Curves USA, NJ) plate reader, at 37°C with continuous shaking. Growth rates were calculated from OD_600_ measurements that were recorded every 5 min for at least 6 hr. This was repeated 6 times for the parental strain PY79, and from 1 to 4 times for all other strains and conditions with PY79 measured in parallel as a growth control.

### TIRF microscopy of Halo-PBP2A

Cultures grown to an OD_600_ below 0.3 were labelled with 10 nM JF549 3, and were concentrated by centrifugation at 6,000 x *g* for 30 sec. The cell pellet was resuspended in growth medium and applied to ethanol-cleaned No. 1.5 coverslips. All cells were imaged under a 2% agarose pad containing growth medium with the top surface exposed to air, in a chamber heated to 37°C. TIRFM and phase-contrast microscopy were performed using a Nikon Eclipse Ti equipped with a Nikon Plan Apo λ 100×/1.45NA objective and a Hamamatsu ORCA-Flash4.0 V2 sCMOS camera. Fluorescence time-lapse images were collected by continuous acquisition with 300 msec exposures. All data are from a single experiment, where cells were induced at different levels, and tracks from >20 cells were used for analysis of each data point.

### Analysis of the density of directionally moving MreB Filaments

Phase images of bacteria were segmented using Morphometrics ^4^, and the width and length of each cell was calculated. Next, the fluorescence time-lapses were analyzed based on the segmentation mask of the phase image (**Figure S3C**). Filament counting was performed in several steps (**Figure S3**). First, kymographs were generated for each row of pixels along the midline of the cell. Next, the kymographs for each row were placed side by side, converting the TIRF time lapse data into a single 2D image (**Figure S3D**). To identify filaments in the kymograph, closed contours were generated in the 2D image (**Figure S3E**). We only selected contours within a given size range (0.04 μm^2^ to 0.17 μm^2^). For these contours, we calculated the total intensity (the sum of the intensities of the pixels in the contour), the centroid, the velocity (calculated from the slope of the major axis line of the contour) (**Figure S3F**), and time (from the centroid). Next, to identify cases where the same MreB filament appears in multiple sequential kymographs, each object in a given kymograph is linked to a corresponding object in the next and previous kymographs based on the above properties of the object (see **Figure S3F** for details). Finally, the counting is verified manually by numbering each filament on the 2D image (**Figure 3A**). To test the performance of the filament counting, we analyzed simulated data with different filament density, velocity, and orientation settings (**Figure S3**). All of the image analyses were performed using MATLAB. Key points in these experiments - specifically, the density of MreB filaments in *E. coli* strains from other labs (**Figure 5**), and the density of MreB filaments in *B. subtilis* (**Figure 3B)** - were repeated twice on independent days. All measurements gave similar mean values and were combined into the final data set.

### Simulation of directionally moving MreB

The Image Correlation Spectroscopy ^5^ MATLAB package was used for the simulation of MreB moving around the cell. The following parameters were set for the MreB simulations: velocity, orientation, filament number and filament length. The default velocity setting is 30 nm/sec and the default orientation is 0, which means MreB moves perpendicular to the central axis. The default filament length is set to 250 nm and each MreB monomer is assumed to be 5 nm. The cell width is set to 1 um and the cell length is set to 4 μm. The pixel size is 65 nm and the time interval is 1 sec, which is the same as the TIRF imaging obtained with our Nikon Eclipse Ti equipped with a Hamamatsu ORCA-Flash4.0 V2 sCMOS camera. The particles are randomly distributed on the surface of the cell. Each simulation data point was repeated 5 times for the counting analysis. To compare the correlated motion approach against particle tracking, we counted the number of tracks observed after tracking the simulated data using the Linear Motion LAP tracker in FIJI ^6^ with TrackMate v3.8 ^7^. The threshold for spot size was 0.195 μm and the intensity threshold was 10 counts. The search radius is 0.085 μm, the link radius is 0.085 μm, and the gap size is 1. All traces longer than 6 frames that had moved more than 250 nm were considered directional motions. Each simulation was repeated 5 times.

### TIRF-SIM imaging of *E. coli* strains

Cells were prepared as described in “Media and culture conditions”. Cells were placed under an LB agarose pad, on a cleaned No.1.5 coverslip, and imaged at 37°C. Live-cell SIM data were acquired as described previously ^8^ on a Zeiss Axio Observer.Z1 inverted microscope outfitted for structured illumination. An Olympus 100×/1.49NA objective was used instead of the Zeiss 1.45NA objective because the slightly larger NA of the Olympus objective gives higher tolerance for placing the excitation beams inside the TIRF annulus. Data was acquired at 1 sec frame rates, with 20 msec exposures from a 488 nm laser for each rotation. TIRF-SIM images were reconstructed as described previously ^8^

### TIRF-SIM imaging of *B. subtilis* strains

Cells were prepared as described in “Media and culture conditions”. Cells were placed under a CH agarose pad in a No. 1.5 glass-bottomed dish (MatTek Corp., MA) for imaging. Images were collected on a DeltaVision OMX SR Blaze system in TIRF mode, using an Edge 5.5 sCMOS camera (PCO AG, Germany) and a 60x objective. 75 msec exposures from a 488 nm diode laser were used for each rotation. Spherical aberration was minimized using immersion oil matching. Raw images were reconstructed using SoftWoRx (GE Healthcare, MA) software.

### Particle tracking of JF549-Halo-PBP2A

Particle tracking was performed using the software package FIJI ^6^ and the TrackMate v3.8 ^7^ plugin. For calculation of particle velocity, the scaling exponent α, and track orientations relative to the midline of the cell, only tracks persisting for 7 frames or longer were used. Particle velocity for each track was calculated from nonlinear least squares fitting using the equation MSD(t) = 4Dt + (vt)^2^, where MSD is mean squared displacement, t is time interval, D is the diffusion coefficient, and v is speed. The maximum time interval used was 80% of the track length. To filter for directionally moving tracks, we discarded those with a velocity lower than 0.01 nm/sec. Tracks were also excluded if the R^2^ for log MSD versus log t was less than 0.95, indicating a poor ability to fit the MSD curve.

### Osmotic shock experiments

Overnight, exponentially growing cultures (as described in “Media and culture conditions”) were diluted into fresh CH medium, grown at 37°C to an OD_600_ of 0.1 to 0.2, then stained by growing in 100 μM Alexa Fluor 488-D-Lysine-NH2 for 1 hr. Without washing, cells were then loaded into a CellASIC microfluidic flow cell (MilliporeSigma, MA) pre-conditioned with media and washed in the chamber via channels 6 and 5. Media in channel 6 was replaced with 5 M NaCl, and the flow cell was resealed and imaged immediately. After collecting images across the whole chip pre-shock, 5 M NaCl was flowed into the chip via channel 6 and imaged immediately. Each “Width” and “Length” data point (**Figure S4B)** is calculated from at least 145 cells (through >1000; see **Table S1**) from multiple fields of view across different areas of the flow cell.

### Alexa Fluor 488-D-Lysine-NH2

Alexa Fluor 488-D-Lysine-NH2 was synthesized as in Lebar et al., 2014. Briefly, Boc-D-Lys(Cbz)-OH (Bachem, Switzerland) was reacted with carbonyldiimidazole (CDI) (MilliporeSigma, MA) in dimethylformamide (DMF) for 1.5 hr, then aqueous ammonia was added and stirred for 6 hr to form the carboxamide Boc-D-Lys(Cbz)-NH2. The Cbz protecting group was removed by catalytic hydrogenation (20% Pd(OH)2/C) in methanol. The product, Boc-D-Lys-NH2, was reacted with CDI in DMF for 1.5 hr, then Alexa Fluor 488 carboxylic acid in DMF was added and reacted for 6 hr to yield Boc-D-Lys(Alexa Fluor 488)-NH2. The Boc protecting group was removed by stirring in neat trifluoroacetic acid (TFA) for 30 min. The reaction was stopped by dropwise addition of TFA solution to ice-cold ether. The precipitate was then HPLC-purified to obtain Alexa Fluor 488-D-Lysine-NH2.

### CRISPRi titration of MreB expression

We used complementarity-based CRISPR knockdown to titrate the MreB expression level in *E. coli*. The degree of MreB-SW-msfGFP repression is controlled by introducing mismatches between the guide RNA and the target DNA ^9^. The repression strength can be tuned by modulating spacer complementarity to msfGFP using different numbers of mismatches. To repress msfGFP using CRISPR knockdown, we placed the dCas9 cassette under the control of a Ptet promoter and different plasmids to target msfGFP. We induced dCas9 at a constant high level with 100 ng/ml of ATc and changed the degree of guide RNA complementarity with different plasmids. For pcrRNA plasmid we use four different guide RNAs with 10, 11, 14, and 20 bp of complementarity. For pAV20 plasmid we use four different guide RNAs with 5, 10, 14, and 20 bp of complementarity. The cells were grown and imaged in LB containing 50 ug/ml kanamycin. This experiment was repeated twice, yielding similar means for each data point. All repeats were then combined into the final data set.

### Protein extraction and labelling

Cell cultures were grown to an OD_600_ of ∼0.4 to 0.6 (cell amounts were normalized by harvesting the equivalent of 3 mL of culture at an OD_600_ of 0.5). Cells were centrifuged at 6,000 × *g* for 30 sec, washed once in 1 mL of ice-cold 20mM Tris-HCl, pH 7.5, and 10 mM EDTA (TE), were resuspended in 100 μL of TE, and were killed by boiling in a water bath for 10 min. All samples were frozen at −80°C overnight (or up to 1 week). Thawed samples were digested with 50 μg of lysozyme (Thermo Fisher, MA) in the presence of 1 mM PMSF, at 37°C for 15 min.

Protein extraction was achieved utilizing a Covaris S220 ultrasonicator (Covaris, MA), under denaturing conditions upon the addition of urea-based Protein Extraction Buffer DF (Covaris, MA), followed by ice-cold methanol/chloroform precipitation. Proteins were digested with trypsin (Promega, WI). Each resulting peptide mixture was labeled with one of a set of up to eleven isotopic tandem mass tags (TMTs) (Thermo Fisher, MA).

### Peptide fractionation and mass spectrometry

Equal amounts of each TMT-labelled sample were combined and fractionated by electrostatic repulsion-hydrophobic interaction chromatography, on an Agilent 1200 HPLC system (Agilent, CA) using a PolyWAX LP 200 × 2.1 mm, 5 μm, 300Â column (PolyLC, MD). Peptides were separated across a 70 min gradient from 0% of “buffer A” (90% acetonitrile, 0.1% acetic acid) to 75% of “buffer B” (30% acetonitrile, 0.1% formic acid), with 20 fractions collected over time. Each fraction was dried in a SpeedVac (Eppendorf, Germany) and resuspended in 0.1% formic acid before injection to a mass spectrometer.

LC-MS/MS was performed on a Thermo Orbitrap Elite (Thermo Fisher, MA) mass spectrometer equipped with a Waters nanoACQUITY HPLC pump (Waters Corp., MA). Peptides were separated on a 150 μm inner diameter microcapillary trapping column packed with ∼3 cm of C18 Reprosil 5 μm, 100 Å resin (Dr. Maisch GmbH, Germany), followed by an analytical column packed with ∼20 cm of Reprosil 1.8 μm, 200 Å resin. Separation was achieved by applying a gradient of 5–27% acetonitrile in 0.1% formic acid, over 90 min at 200 nl/min. Electrospray ionization was achieved by applying a voltage of 2 kV using a home-made electrode junction at the end of the microcapillary column and sprayed from fused-silica PicoTips (New Objective, MA). The Orbitrap instrument was operated in data-dependent mode for the mass spectrometry methods. The mass spectrometry survey scan was performed in the Orbitrap in the range of 410 –1,800 m/z at a resolution of 12 × 10^4^, followed by the selection of the twenty most intense ions for HCD-MS2 fragmentation using a precursor isolation width window of 2 m/z, AGC setting of 50,000, and a maximum ion accumulation of 200 msec. Singly-charged ion species were not subjected to HCD fragmentation. Normalized collision energy was set to 37 V and an activation time of 1 msec. Ions in a 10 ppm m/z window around ions selected for MS2 were excluded from further selection for fragmentation for 60 sec.

Mass spectrometry analysis: Raw data were submitted for analysis in Proteome Discoverer 2.1.0.81 (Thermo Scientific) software. Assignment of MS/MS spectra was performed using the Sequest HT algorithm by searching the data against a protein sequence database that including all entries from *B. subtilis* (UniProt proteome ID UP000018540) and other known contaminants such as human keratins and common lab contaminants. Sequest HT searches were performed using a 15 ppm precursor ion tolerance and required each peptide’s N- and C-termini to adhere with trypsin protease specificity, while allowing up to two missed cleavages. TMT tags on peptide N-termini and lysine residues (+229.162932 Da) were set as static modifications while methionine oxidation (+15.99492 Da) was set as a variable modification. An MS2 spectra assignment false discovery rate (FDR) of 1% on protein level was achieved by applying the target-decoy database search. Filtering was performed using Percolator (64-bit version ^10^). For quantification, a 0.02 m/z window centered on the theoretical m/z value of each of the TMT reporter ions and the intensity of the signal closest to the theoretical m/z value was recorded. Reporter ion intensities were exported in a results file of the Proteome Discoverer 2.1 search engine in Microsoft Excel format. The total signal intensity across all peptides quantified was summed for each TMT channel, and all intensity values were adjusted to account for potentially uneven TMT-labelling and/or sample handling variance for each labelled channel. For our final relative protein quantitation analysis, all contaminants from the database search were removed from the results, and only the remaining *B. subtilis* proteins were used to re-normalize all protein abundances.

### Strain construction

**bMD277** containing *amyE::Pxyl-mreBCD, minCD (B. megaterium)::erm* was generated upon transformation of PY79 with a five-piece Gibson assembly reaction^11^, that contained the following PCR products. (1) A 1228 bp fragment containing sequence upstream of the *amyE* locus (amplified from PY79 genomic DNA using primers oMD191 and oMD108); (2) the 1673 bp erythromycin-resistance cassette *loxP-erm-loxP* (amplified from pWX467 [gift of D. Rudner] using primers oJM028 and oJM029); (3) a 1532 bp fragment containing the *xylR* gene, and the *PxylA* promoter with an optimized ribosomal binding sequence (amplified from pDR150 [gift of D. Rudner] using primers oMD73 and oMD226); (4) a 4106 bp fragment containing the *mreB, mreC, mreD, minC* and *minD* alleles of the *B. megaterium mreB* operon (amplified from strain QMB 1551 (ATCC 12872) genomic DNA using primers oMD227 and oMD228); and (5) a 1216 bp fragment containing the *amyE* terminator, and sequence downstream of the *amyE* locus (amplified from PY79 genomic DNA using primers oMD196 and oMD197).

**bMD465** harboring *amyE::Pxyl-mreBCD, minCD (B. megaterium)::erm, mreBCD, minC, D::spec (B. megaterium)* was generated upon transformation of bMD277 with a four-piece Gibson assembly reaction, that contained the following PCR products. (1) A 1275 bp fragment containing sequence upstream of the *mreB* gene (amplified from PY79 genomic DNA using primers oMD96 and oMD308); (2) a 4121 bp fragment containing the *mreB, mreC, mreD, minC* and *minD* alleles of the *B. megaterium mreB* operon (amplified from strain QMB 1551 (ATCC 12872) genomic DNA using primers oMD313 and oMD314); (3) the 1274 bp spectinomycin-resistance cassette *loxP-spec-loxP* (amplified from pWX466 [gift of D. Rudner] using primers oJM028 and oJM029); and (4) a 1152 bp fragment containing sequence downstream of the *minD* gene (amplified from PY79 genomic DNA using primers oMD300 and oMD315).

**bMK355** harboring *amyE::Pxyl-mreBCD::erm* was built identical to bMD277, except that PCR product (4) was instead a 2493 bp fragment containing the *mreB*, *mreC*, and *mreD* alleles of the *B. subtilis mreB* operon (amplified from PY79 genomic DNA using primers oMD334 and oMK221).

**bMD543** containing *amyE::Pxyl-mreBCD::erm, ΔmreBCD, ΔminCD::cat* was generated upon transformation of bMK355 with a three-piece Gibson assembly reaction, that contained the following PCR products. (1) A 1305 bp fragment containing sequence upstream of the *mreB* gene (amplified from PY79 genomic DNA using primers oMD96 and oMD298); (2) the 1139 bp chloramphenicol-resistance cassette *loxP-cat-loxP* (amplified from pWX465 [gift of D. Rudner] using primers oJM028 and oJM029); and (3) a 1215 bp fragment containing sequence downstream of the *minD* gene (amplified from PY79 genomic DNA using primers oMD299 and oMD300).

**bMD545** containing *amyE::Pxyl-mreBCD::erm, ΔmreBCD, PmreB-minC, D::spec* was generated upon transformation of bMD543 with a four-piece Gibson assembly reaction, that contained the following PCR products. (1) A 1305 bp fragment containing sequence upstream of the *mreB* gene (amplified from PY79 genomic DNA using primers oMD96 and oMD379); (2) a 1582 bp fragment containing the *minC* and *minD* genes, and the *minD* terminator of the *mreB* operon (amplified from PY79 genomic DNA using primers oMD380 and oMD381); (3) the 1274 bp spectinomycin-resistance cassette *loxP-spec-loxP* (amplified from pWX466 using primers oJM028 and oJM029); and (4) a 1215 bp fragment containing sequence downstream of the *minD* gene (amplified from PY79 genomic DNA using primers oMD300 and oMD382).

**bMD599** harboring *ΔponA::kan* was generated upon transformation of PY79 with a three-piece Gibson assembly reaction, that contained the following PCR products. (1) A 1329 bp fragment containing sequence upstream of the *ponA* gene (amplified from PY79 genomic DNA using primers oMK001 and oMK002); (2) the 1577 bp kanamycin-resistance cassette *loxP-kan-loxP* (amplified from pWX470 [gift of D. Rudner] using primers oJM028 and oJM029); and (3) a 1313 bp fragment containing sequence downstream of the *ponA* gene (amplified from PY79 genomic DNA using primers oMK005 and oMK006).

**bMK005** containing *ΔponA::cat* was built identical to bMD599, except that PCR product (2) was instead the 1139 bp chloramphenicol-resistance cassette *loxP-cat-loxP* (amplified from pWX465 using primers oJM028 and oJM029).

**bMD554** containing *yhdG::Phyperspank-ponA::cat* was generated upon transformation of PY79 with a five-piece Gibson assembly reaction, that contained the following PCR products. (1) A 1219 bp fragment containing sequence upstream of the *yhdH* gene (amplified from PY79 genomic DNA using primers oMD328 and oMD367), (2) the 1139 bp chloramphenicol-resistance cassette *loxP-cat-loxP* (amplified from pWX465 using primers oJM028 and oJM029); (3) a 1895 bp fragment containing the *lacI* gene, and the *Phyperspank* promoter with an optimized ribosomal binding sequence (amplified from pDR111 [gift of D. Rudner] using primers oMD232 and oMD234); (4) a 2799 bp fragment containing the *ponA* coding region (amplified from PY79 genomic DNA using primers oMD365 and oMK370); and (5) a 1252 bp fragment containing the *yhdG* terminator, and sequence downstream of the *yhdG* gene (amplified from PY79 genomic DNA using primers oMD371 and oMD372).

**bMD586** harboring *yhdG::Phyperspank-ponA::cat*, *ΔponA::kan* was generated upon transformation of bMD554 with genomic DNA from bMD599.

**bMD594** harboring *yhdG::Pspank-ponA::cat* was built identical to bMD554, except that PCR product (1) was instead a 1287 bp fragment made with primers oMD328 and oMD329; and PCR product (3) containing the *Pspank* promoter was instead amplified from pDR110 (gift of D. Rudner).

**bMD598** containing *yhdG::Pspank-ponA::cat*, *ΔponA::kan* was generated upon transformation of bMD594 with genomic DNA from bMD599.

**bMD619** containing *amyE::Pxyl-mreBCD::erm, ΔmreBCD, PmreB-minCD::spec*, *yhdG::Pspank-ponA::cat* was generated upon transformation of bMD545 with genomic DNA from bMD594.

**bMD620** containing *amyE::Pxyl-mreBCD::erm, ΔmreBCD, PmreB-minCD::spec*, *yhdG::Pspank-ponA::cat*, *ΔponA::kan* was generated upon transformation of bMD619 with genomic DNA from bMD599.

**bMD622** harboring *amyE::Pxyl-mreBCD::erm*, *yhdG::Pspank-ponA::cat*, *ΔponA::kan* was generated upon transformation of bMD598 with genomic DNA from bMK355.

**bMD556** harboring *yhdG::Phyperspank-rodA::cat* was built identical to bMD554, except that PCR product (4) was instead a 1236 bp fragment containing the *rodA* coding region (amplified from PY79 genomic DNA using primers oMD364 and oMK369).

**bMD580** containing *yhdG::Phyperspank-rodA::cat*, *ΔrodA::kan* was generated upon transformation of bMD556 with a three-piece Gibson assembly reaction, that contained the following PCR products. (1) A 1265 bp fragment containing sequence upstream of the *rodA* gene (amplified from PY79 genomic DNA using primers oMD388 and oMD389); (2) the 1577 bp kanamycin-resistance cassette *loxP-kan-loxP* (amplified from pWX470 using primers oJM028 and oJM029); and (3) a 1258 bp fragment containing sequence downstream of the *rodA* gene (amplified from PY79 genomic DNA using primers oMD386 and oMD387).

**bMD592** harboring *Pxyl-rodA::erm* was generated upon transformation of PY79 with a four-piece Gibson assembly reaction, that contained the following PCR products. (1) A 1265 bp fragment containing sequence upstream of the *rodA* gene (amplified from PY79 genomic DNA using primers oMD388 and oMD389); (2) the 1673 bp erythromycin-resistance cassette *loxP-erm-loxP* (amplified from pWX467 using primers oJM028 and oJM029); (3) a 1532 bp fragment containing the *xylR* gene, and the *PxylA* promoter with an optimized ribosomal binding sequence (amplified from pDR150 using primers oMD73 and oMD226); and (4) a 1271 bp fragment containing the *rodA* coding region (amplified from PY79 genomic DNA using primers oMD394 and oMD395).

**bMD627** containing *Pxyl-rodA::erm*, *ΔpbpH::spec* was generated upon transformation of bMD592 with genomic DNA from bDR2487.

**bMD563** containing *yhdG::Phyperspank-pbpA::cat* was built identical to bMD554, except that PCR product (4) was instead a 2204 bp fragment containing the *pbpA* coding region (amplified from PY79 genomic DNA using primers oMD316 and oMK368).

**bMD573** harboring *yhdG::Phyperspank-pbpA::cat*, *ΔpbpH::spec* was generated upon transformation of bMD563 with genomic DNA from bRB776.

**bMD574** harboring *yhdG::Phyperspank-pbpA::cat*, *ΔpbpA::erm*, *ΔpbpH::spec* was generated upon transformation of bMD573 with genomic DNA from bRB776.

**bMD590** containing *Pxyl-pbpA::erm* was generated upon transformation of PY79 with a four-piece Gibson assembly reaction, that contained the following PCR products. (1) A 1233 bp fragment containing sequence upstream of the *pbpA* gene (amplified from PY79 genomic DNA using primers oMD75 and oMD125); (2) the 1673 bp erythromycin-resistance cassette *loxP-erm-loxP* (amplified from pWX467 using primers oJM028 and oJM029); (3) a 1537 bp fragment containing the *xylR* gene, and the *PxylA* promoter containing the *pbpA* ribosomal binding sequence (amplified from pDR150 using primers oMD73 and oMD72); and (4) a 1230 bp fragment containing the *pbpA* coding region (amplified from PY79 genomic DNA using primers oMD393 and oMD68).

**bMD597** harboring *Pxyl-pbpA::erm*, *ΔpbpH::spec* was generated upon transformation of bMD590 with genomic DNA from bDR2487.

**bMD557** containing *amyE::Pxyl-mreBCD::erm*, *yhdG::Phyperspank-rodA::cat* was generated upon transformation of bMK355 with genomic DNA from bMD556.

**bMD571** harboring *amyE::Pxyl-mreBCD::erm*, *yhdG::Phyperspank-pbpA::cat* was generated upon transformation of bMK355 with genomic DNA from bMD563.

**bMD631** containing *yhdG::Pspank-pbpA::phleo, ΔpbpH::spec, ΔpbpA::cat, Pxyl-rodA::erm* was generated upon transformation of bRB773 with genomic DNA from bMD592.

**bYS201** harboring *HaloTag-11aa-pbpA::cat* was generated upon transformation of PY79 with a four-piece Gibson assembly reaction, that contained the following PCR products. (1) A 1233 bp fragment containing sequence upstream of the *pbpA* gene (amplified from PY79 genomic DNA using primers oMD75 and oMD125); (2) the 1139 bp chloramphenicol-resistance cassette *loxP-cat-loxP* (amplified from pWX465 using primers oJM028 and oJM029); (3) a 962 bp fragment containing the native *pbpA* promoter, and *B. subtilis*-optimized coding sequence of the HaloTag protein (amplified from double-stranded DNA custom-ordered from DNA 2.0/ATUM [Newark, CA] using primers oYS598 and oYS599); and (4) a 1229 bp fragment containing the *pbpA* coding region (amplified from PY79 genomic DNA using primers oMD68 and oMD69).

**bMK385** harboring *amyE::Pxyl-mreBCD::erm, ΔmreBCD, PmreB-minCD::spec*, *HALO-11aa-pbpA::cat* was generated upon transformation of bMD545 with genomic DNA from bYS201.

**bYS19** containing *mreB-SW-msfGFP* was generated upon transformation of bMD88 ^12^ with a three-piece Gibson assembly reaction that contained the following PCR products. (1) A 898 bp PCR fragment containing sequence upstream of the *mreB* gene (amplified from PY79 genomic DNA using primers oMD134 and oYS34); (2) a 774 bp fragment containing *B. subtilis*-optimized coding sequence of the msfGFP fluorescent protein (amplified from plasmid DNA custom-ordered from DNA 2.0/ATUM [Newark, CA] using primers oYS35 and oYS36); and (3) a 1629 bp fragment containing sequence downstream of the *mreB* gene (amplified from PY79 genomic DNA using primers oYS37 and oMD116). This strain is markerless, and selection was in the presence of 0.5% glucose for colonies that no longer required xylose for viability.

**bYS977** harboring *amyE::Pxyl-mreB-SW-msfGFP, mreCD::erm*, was built identical to bMD277, except that PCR product (4) was instead a 3234 bp fragment containing *mreB-SW-msfGFP*, *mreC*, and *mreD*, amplified from bYS19 genomic DNA using primers oMD334 and oMK221).

**bYS979** containing *amyE::Pxyl-mreB-SW-msfGFP, mreCD::erm, ΔmreBCD, ΔminCD::cat* was generated upon transformation of bYS977 with genomic DNA from bMD543.

**bYS981** harboring *amyE::Pxyl-mreB-SW-msfGFP, mreCD::erm, ΔmreBCD, PmreB-minC, D::spec* was generated upon transformation of bYS979 with genomic DNA from bMD545.

**AV88** containing *186::Ptet-dCas9, mreB-SW-msfGFP* was obtained by transduction of strain LC69 with a P1 phage lysate obtained from strain NO50, and subsequent excision of the kanamycin resistance cassette by transient expression of a flippase. To make the plasmids expressing single-guide RNAs, the psgRNA plasmid was amplified three times with primers pairs V162/V165, V163/V166 and V164/V167. The three generated fragments were assembled together by Gibson assembly to obtain the pAV20 vector. To insert the anti-GFP single-guide RNAs into this vector, we annealed and phosphorylated the following pairs of oligonucleotides: V275/V283 for G5; V274/V282 for G10; V273/V281 for G14; and V272/V280 for G20. These were then inserted in pAV20 by restriction cloning with *Bsa*I. As pAV20 carries two sgRNA insertion sites, a non-targeting oligonucleotide pair (V279/V287) was inserted into a second site during the same reaction.

## Statistics

All P-values are reported in figures. P-values were calculated in GraphPad Prism using the Mann-Whitney test with a two-tailed P value. Means, SD, and N for all data points are reported in table S1. Fits to data in Figures 3B and 5A were done using linear regression in GraphPad Prism 7.0. Replicates of experiments are reported in Materials and Methods.

## Data Availability

All datasets generated during and/or analyzed during the current study are available from the corresponding author on reasonable request. All proteomic and other data are available at https://garnerlab.fas.harvard.edu/Dion2018/

## Software

All particle tracking was done with the Trackmate plugin within Fiji, then analyzed using code available at: https://bitbucket.org/garnerlab/hussain-2017-elife. Filament density calculations and filament simulations were done with custom code available at: https://bitbucket.org/garnerlab/dion-2018/src/

---

**Table S1.** – Data summary and statistics.

**A - Figure 1.**

A - Figure 1.
| Panel | Strain | condition. | parameter | Value | SD | N |
| --- | --- | --- | --- | --- | --- | --- |
| B | PY79 |  | width(nm) | 754.70 | 81.19 | 466 |
| B | *B. megaterium* |  | width(nm) | 1377.88 | 98.09 | 337 |
| B | bMD465 | 1% glucose | width(nm) | 807.52 | 61.24 | 267 |
| B | bMD465 | 30 mM xylose | width(nm) | 771.20 | 104.12 | 135 |
| C | bMD545 | 0.1 mM xylose | width(nm) | 1533.00 | 247.40 | 501 |
| C | bMD545 | 0.5 mM xylose | width(nm) | 1179.00 | 165.50 | 466 |
| C | bMD545 | 1 mM xylose | width(nm) | 1059.00 | 145.70 | 395 |
| C | bMD545 | 1.75 mM xylose | width(nm) | 930.80 | 83.53 | 282 |
| C | bMD545 | 2.5 mM xylose | width(nm) | 807.80 | 90.69 | 461 |
| C | bMD545 | 5 mM xylose | width(nm) | 735.00 | 80.10 | 269 |
| C | bMD545 | 30 mM xylose | width(nm) | 722.10 | 60.64 | 619 |
| C | bMK355 | 30 mM xylose | width(nm) | 696.50 | 85.40 | 276 |
| D | bMD599 |  | width(nm) | 575.90 | 97.00 | 174 |
| D | bMD598 | 0 M IPTG | width(nm) | 580.60 | 60.57 | 148 |
| D | bMD598 | 1 μM IPTG | width(nm) | 601.00 | 68.93 | 141 |
| D | bMD598 | 10 μM IPTG | width(nm) | 651.40 | 74.63 | 742 |
| D | bMD598 | 25 μM IPTG | width(nm) | 767.80 | 91.60 | 309 |
| D | bMD598 | 100 μM IPTG | width(nm) | 825.00 | 78.83 | 361 |
| D | bMD598 | 1000 μM IPTG | width(nm) | 868.70 | 78.53 | 654 |
| D | bMD586 | 25 μM IPTG | width(nm) | 935.30 | 106.00 | 332 |
| D | bMD586 | 50 μM IPTG | width(nm) | 1031.30 | 97.14 | 381 |
| D | bMD586 | 100 μM IPTG | width(nm) | 1149.45 | 119.40 | 133 |
| D | bMD586 | 1000 μM IPTG | width(nm) | 1283 | 137.8 | 619 |
| D | bMD554 | 1000 μM IPTG | width(nm) | 1519 | 169.4 | 276 |
| E | bMD620 | 0.1 mM xylose, 0 M IPTG | width(nm) | 786.00 | 159.30 | 452 |
| E | bMD620 | 0.5 mM xylose, 1 μM IPTG | width(nm) | 744.50 | 137.90 | 265 |
| E | bMD620 | 2.5 mM xylose, 10 μM IPTG | width(nm) | 754.40 | 76.76 | 522 |
| E | bMD620 | 5 mM xylose, 25 μM IPTG | width(nm) | 749.00 | 73.91 | 288 |
| E | bMD620 | 30 mM xylose, 100 μM IPTG | width(nm) | 784.80 | 76.63 | 589 |
| E | bMD622 | 30 mM xylose, 1000 μM IPTG | width(nm) | 773.30 | 58.09 | 665 |
| F | PY79 | LB | width(nm) | 770.50 | 44.09 | 590 |
| F | PY79 | CH | width(nm) | 754.70 | 81.19 | 466 |
| F | PY79 | S750 glucose | width(nm) | 767.49 | 78.52 | 389 |
| F | PY79 | S750 Glycerol | width(nm) | 756.90 | 70.21 | 596 |
| F | PY79 | S750 Sucrose | width(nm) | 761.51 | 63.79 | 360 |

**B - Figure 2**

**B - Figure 2**
| Panel | Strain | Conditions | parameter | Value | SD | N |
| --- | --- | --- | --- | --- | --- | --- |
| A | bMD592 | 0.1mM xyl | width(nm) | 1113 | 221.8 | 387 |
| A | bMD592 | 0.5mM xyl | width(nm) | 959.8 | 156.6 | 246 |
| A | bMD592 | 1mM xyl | width(nm) | 753.2 | 106.6 | 370 |
| A | bMD592 | 30mM xyl | width(nm) | 964.2 | 77.79 | 466 |
| A | bMD580 | 1mM IPTG | width(nm) | 1133 | 110.8 | 555 |
| A | bMD556 | 1mM IPTG | width(nm) | 1323 | 155.5 | 788 |
| A | bMD598 | 0.1mM xyl | width(nm) | 1081 | 206.5 | 455 |
| A | bMD599 | 0.5mM xyl | width(nm) | 927.6 | 117.8 | 442 |
| A | bMD600 | 1mM xyl | width(nm) | 749.3 | 77.61 | 135 |
| A | bMD601 | 30mM xyl | width(nm) | 782 | 52.83 | 91 |
| A | bMD602 | 1mM IPTG | width(nm) | 790.1 | 66.96 | 288 |
| A | bMD603 | 1mM IPTG | width(nm) | 795.1 | 59.96 | 443 |
| B | bMD627 | 0.025mM xyl | width(nm) | 1381 | 334.7 | 92 |
| B | bMD627 | 1mM xyl | width(nm) | 756.4 | 52.33 | 983 |
| B | bMD627 | 30mM xyl | width(nm) | 793.4 | 62.71 | 414 |
| B | bMD631 | 0.25μM xyl, 15μM IPTG | width(nm) | 1346 | 371.4 | 79 |
| B | bMD631 | 1mM xyl, 15μM IPTG | width(nm) | 757.9 | 55.22 | 441 |
| B | bMD631 | 30mM xyl 15μM IPTG | width(nm) | 966.1 | 80.25 | 503 |
| B | bMD631 | 30mM xyl 20μM IPTG | width(nm) | 902 | 106.4 | 262 |
| C | bMD557 | 30mM xyl, 40μM IPTG | width(nm) | 987.1 | 87.58 | 230 |
| C | bMD557 | 30mM xyl 100μM IPTG | width(nm) | 856.5 | 121.3 | 266 |
| D | bMD620 | 0.1mM xyl 0 IPTG | Surface growth | 0.755 | 0.1681 | 183 |
| D | bMD620 | 0.5mM xyl 1μM IPTG | Surface growth | 1.025 | 0.171 | 63 |
| D | bMD620 | 2.5mM xyl 10μM IPTG | Surface growth | 1.146 | 0.1108 | 57 |
| D | bMD620 | 5mM xyl 25μM IPTG | Surface growth | 1.174 | 0.0692 | 78 |
| D | bMD620 | 30mM xyl 100μM IPTG | Surface growth | 1.282 | 0.1895 | 46 |
| D | bMD622 | 30mM xyl 1mM IPTG | Surface growth | 1.194 | 0.1063 | 54 |
| D | bMD620 | 0.1mM xyl 25μM IPTG | Surface growth | 0.816 | 0.2653 | 17 |
| D | bMD620 | 0.5mM xyl 25μM IPTG | Surface growth | 1.203 | 0.138 | 111 |
| D | bMD620 | 1mM xyl 25μM IPTG | Surface growth | 1.182 | 0.174 | 59 |
| D | bMD620 | 2.5mM xyl 25μM IPTG | Surface growth | 1.188 | 0.1229 | 121 |
| D | bMD620 | 5mM xyl 25μM IPTG | Surface growth | 1.174 | 0.0691 | 78 |
| D | bMD620 | 30mM xyl 25μM IPTG | Surface growth | 1.326 | 0.1969 | 50 |
| D | bMD620 | 5mM xyl 1μM IPTG | Surface growth | 1.274 | 0.182 | 78 |
| D | bMD620 | 5mM xyl 10μM IPTG | Surface growth | 1.256 | 0.1302 | 140 |
| D | bMD620 | 5mM xyl 25μM IPTG | Surface growth | 1.174 | 0.0692 | 78 |
| D | bMD620 | 5mM xyl 50μM IPTG | Surface growth | 1.288 | 0.1677 | 486 |
| D | bMD620 | 5mM xyl 100μM IPTG | Surface growth | 1.353 | 0.1076 | 123 |
| D | bMD599 |  | Surface growth | 1.263 | 0.1487 | 84 |
| D | bAM268 |  | Surface growth | 1.147 | 0.1432 | 177 |
| D | bAM288 | 25μM IPTG | Surface growth | 1.107 | 0.1121 | 51 |
| D | bMD631 | 1mM xyl 15μM IPTG | Surface growth | 1.398 | 0.1488 | 187 |
| D | bMD631 | 30mM xyl 15μM IPTG | Surface growth | 1.327 | 0.0789 | 49 |
| D | bMD631 | 30mM xyl 40μM IPTG | Surface growth | 1.173 | 0.1377 | 47 |
| D | PY79 |  | Surface growth | 1.223 | 0.1527 | 84 |
| D | bMD620 | 0.1mM xyl 0 IPTG | Bulk growth | 1.49 | 0.1 | 5 |
| D | bMD620 | 0.5mM xyl 1μM IPTG | Bulk growth | 1.76 | 0.09 | 10 |
| D | bMD620 | 2.5mM xyl 10μM IPTG | Bulk growth | 1.85 | 0.17 | 9 |
| D | bMD620 | 5mM xyl 25μM IPTG | Bulk growth | 1.85 | 0.07 | 5 |
| D | bMD620 | 30mM xyl 100μM IPTG | Bulk growth | 1.89 | 0.05 | 5 |
| D | bMD622 | 30mM xyl 1mM IPTG | Bulk growth | 2 | 0.03 | 5 |
| D | bMD620 | 0.1mM xyl 25μM IPTG | Bulk growth | 1.277 | 0.07 | 5 |
| D | bMD620 | 0.5mM xyl 25μM IPTG | Bulk growth | 1.98 | 0.05 | 5 |
| D | bMD620 | 1mM xyl 25μM IPTG | Bulk growth | 1.88 | 0.04 | 5 |
| D | bMD620 | 2.5mM xyl 25μM IPTG | Bulk growth | 1.85 | 0.03 | 5 |
| D | bMD620 | 5mM xyl 25μM IPTG | Bulk growth | 2.05 | 0.13 | 10 |
| D | bMD620 | 30mM xyl 25μM IPTG | Bulk growth | 1.97 | 0.03 | 5 |
| D | bMD620 | 5mM xyl 1μM IPTG | Bulk growth | 1.84 | 0.18 | 10 |
| D | bMD620 | 5mM xyl 10μM IPTG | Bulk growth | 1.96 | 0.14 | 9 |
| D | bMD620 | 5mM xyl 25μM IPTG | Bulk growth | 1.76 | 0.05 | 5 |
| D | bMD620 | 5mM xyl 50μM IPTG | Bulk growth | 1.83 | 0.03 | 5 |
| D | bMD620 | 5mM xyl 100μM IPTG | Bulk growth | 1.86 | 0.05 | 5 |
| D | bMD599 |  | Bulk growth | 1.19 | 0.04 | 10 |
| D | bAM268 |  | Bulk growth | 1.17 | 0.03 | 5 |
| D | bAM288 | 25μM IPTG | Bulk growth | 1.7 | 0.02 | 5 |
| D | PY79 |  | Bulk growth | 2 | 0.13 | 26 |

**C - Figure 3**

**C - Figure 3**
| Panel | Strain | Conditions | Width (um) | Width SD | (N) | Filament density | Density SD |
| --- | --- | --- | --- | --- | --- | --- | --- |
| B | YS981 | 0.1mM Xyl | 1.3826 | 0.2 | 336 | 0.29 | 0.28 |
| B | YS981 | 0.25mM Xyl | 1.2808 | 0.2894 | 214 | 0.37 | 0.35 |
| B | YS981 | 0.5mM Xyl | 1.2557 | 0.1674 | 130 | 0.55 | 0.42 |
| B | YS981 | 1mM Xyl | 1.0093 | 0.1128 | 113 | 1.49 | 0.85 |
| B | YS981 | 1.75mM Xyl | 1.0661 | 0.0969 | 116 | 1.36 | 0.58 |
| B | YS981 | 2.5mM Xyl | 1.0469 | 0.1315 | 107 | 1.64 | 0.48 |
| B | YS981 | 5mM Xyl | 1.0051 | 0.0985 | 92 | 2.14 | 0.85 |
| B | YS981 | 30mM Xyl | 1.0124 | 0.0731 | 190 | 2.12 | 0.83 |

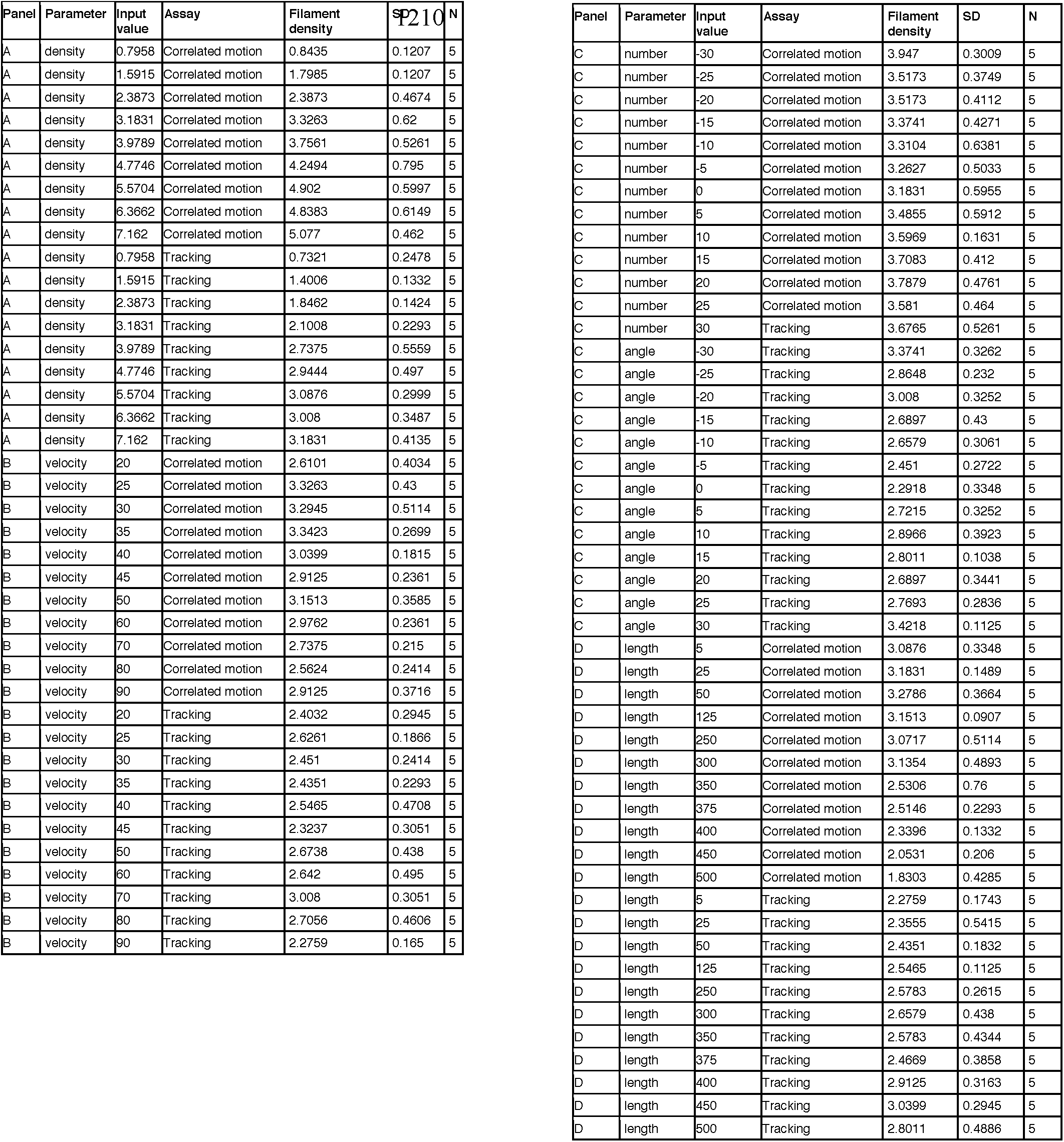
**D - Figure S3 – Simulation**

**E - Figure 4, S4**

**E - Figure 4, S4**
| Panel | Strain | Conditions | Parameter | Value | SD | (N) |
| --- | --- | --- | --- | --- | --- | --- |
| B, D | bMD620 | 30/0.001 | Width | 1.3826 | 0.2 | 93 |
| B, D | bMD620 | 5/0.025 | Width | 1.2808 | 0.2894 | 214 |
| B, D | bMD620 | 0.5/100 | Width | 1.2557 | 0.1674 | 130 |
| D, E | bMD620 | 30/0.001 | Retardance | 0.3937 | 0.0632 | 93 |
| D, E | bMD620 | 5/0.025 | Retardance | 0.3676 | 0.0468 | 91 |
| D, E | bMD620 | 0.5/100 | Retardance | 0.2824 | 0.0583 | 137 |
| S4A | bMD620 | 30/0.001 | Cell width (TEM) | 512.38 | 45.5616 | 16 |
| S4A | bMD620 | 5/0.025 | Cell width (TEM) | 705.71 | 51.6678 | 35 |
| S4A | bMD620 | 0.5/100 | Cell width (TEM) | 1138.8 | 229.043 | 58 |
| S4A | PY79 |  | Cell width (TEM) | 748.08 | 31.5936 | 32 |
| S4A | bMD620 | 30/0.001 | Cell wall width (TEM) | 14 | 3.2609 | 16 |
| S4A | bMD620 | 5/0.025 | Cell wall width (TEM) | 15.34 | 1.6439 | 35 |
| S4A | bMD620 | 0.5/100 | Cell wall width (TEM) | 19.51 | 3.7475 | 58 |
| S4A | PY79 |  | Cell wall width (TEM) | 15.11 | 1.3955 | 32 |
| H, S4B, S4C | bMD620 | 5/0.025 preshock | Sacculus length | 4470 | 1138 | 446 |
| H, S4B, S4C | bMD620 | 5/0.025 preshock | Sacculus Width | 782.6 | 53.03 | 446 |
| H, S4B, S4C | bMD620 | 5/0.025 shocked | Sacculus length | 3120 | 973.5 | 309 |
| H, S4B, S4C | bMD620 | 5/0.025 shocked | Sacculus Width | 676.6 | 41.38 | 309 |
| H, S4B, S4C | bMD620 | 0.5/100 preshock | Sacculus length | 4261 | 1063 | 145 |
| H, S4B, S4C | bMD620 | 0.5/100 preshock | Sacculus Width | 1283 | 199.2 | 145 |
| H, S4B, S4C | bMD620 | 0.5/100 shocked | Sacculus length | 3386 | 812.6 | 146 |
| H, S4B, S4C | bMD620 | 0.5/100 shocked | Sacculus Width | 1085 | 164.9 | 146 |
| H, S4B, S4C | bMD620 | 30/0.001 preshock | Sacculus length | 3789 | 1535 | 1112 |
| H, S4B, S4C | bMD620 | 30/0.001 preshock | Sacculus Width | 586.4 | 43.09 | 1112 |
| H, S4B, S4C | bMD620 | 30/0.001 shocked | Sacculus length | 2315 | 919.6 | 408 |
| H, S4B, S4C | bMD620 | 30/0.001 shocked | Sacculus Width | 527.6 | 59.77 | 408 |

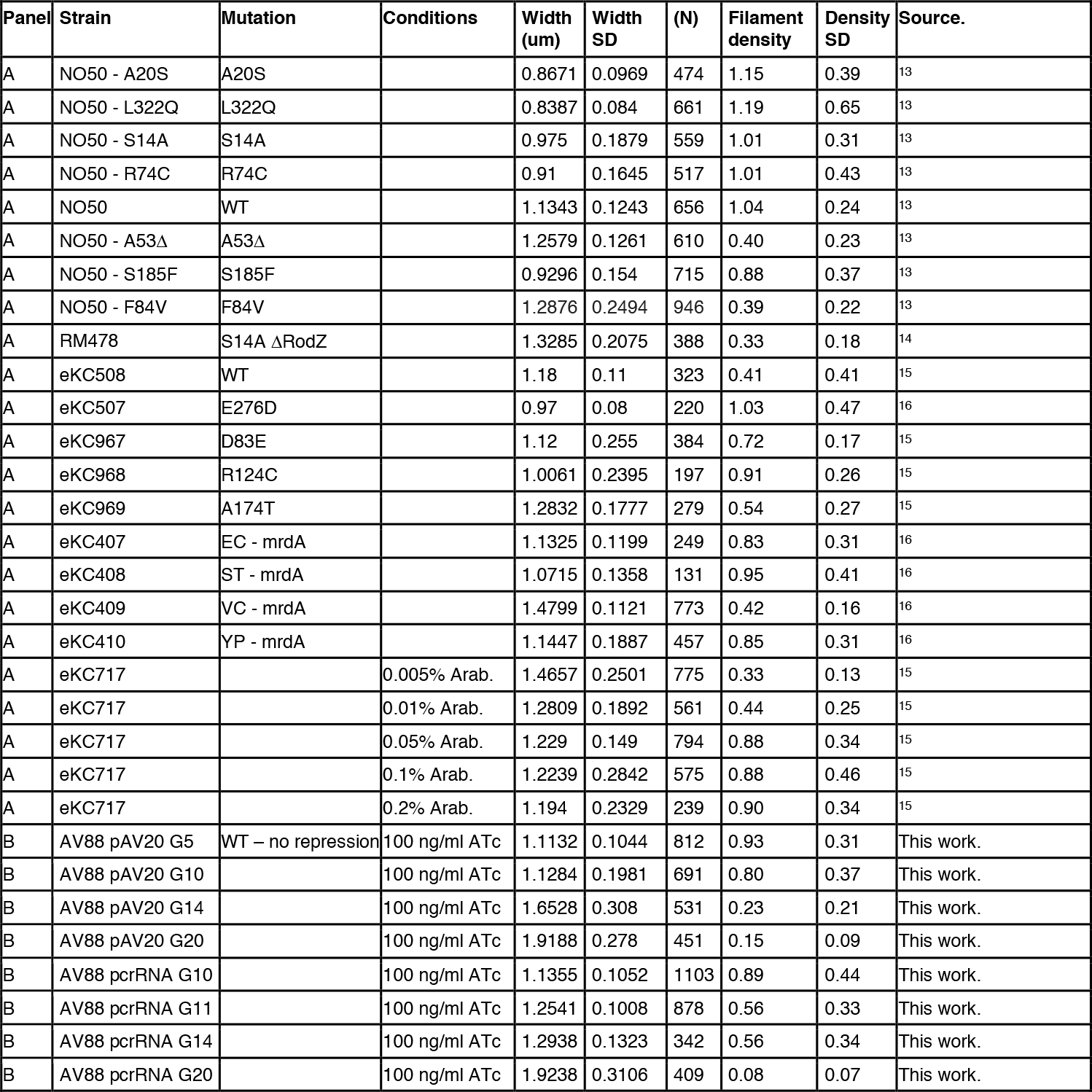
**F - Figure 5.**

F - Figure 5.
| Panel | Strain | Mutation | Conditions | Width (um) | Width SD | (N) | Filament density | Density SD | Source. |
| --- | --- | --- | --- | --- | --- | --- | --- | --- | --- |
| A | NO50 - A20S | A20S |  | 0.8671 | 0.0969 | 474 | 1.15 | 0.39 | <sup>13</sup> |
| A | NO50 - L322Q | L322Q |  | 0.8387 | 0.084 | 661 | 1.19 | 0.65 | <sup>13</sup> |
| A | NO50 - S14A | S14A |  | 0.975 | 0.1879 | 559 | 1.01 | 0.31 | <sup>13</sup> |
| A | NO50 - R74C | R74C |  | 0.91 | 0.1645 | 517 | 1.01 | 0.43 | <sup>13</sup> |
| A | NO50 | WT |  | 1.1343 | 0.1243 | 656 | 1.04 | 0.24 | <sup>13</sup> |
| A | NO50 - A53Δ | A53Δ |  | 1.2579 | 0.1261 | 610 | 0.40 | 0.23 | <sup>13</sup> |
| A | NO50 - S185F | S185F |  | 0.9296 | 0.154 | 715 | 0.88 | 0.37 | <sup>13</sup> |
| A | NO50 - F84V | F84V |  | 1.2876 | 0.2494 | 946 | 0.39 | 0.22 | <sup>13</sup> |
| A | RM478 | S14A ΔRodZ |  | 1.3285 | 0.2075 | 388 | 0.33 | 0.18 | <sup>14</sup> |
| A | eKC508 | WT |  | 1.18 | 0.11 | 323 | 0.41 | 0.41 | <sup>15</sup> |
| A | eKC507 | E276D |  | 0.97 | 0.08 | 220 | 1.03 | 0.47 | <sup>16</sup> |
| A | eKC967 | D83E |  | 1.12 | 0.255 | 384 | 0.72 | 0.17 | <sup>15</sup> |
| A | eKC968 | R124C |  | 1.0061 | 0.2395 | 197 | 0.91 | 0.26 | <sup>15</sup> |
| A | eKC969 | A174T |  | 1.2832 | 0.1777 | 279 | 0.54 | 0.27 | <sup>15</sup> |
| A | eKC407 | EC - mrdA |  | 1.1325 | 0.1199 | 249 | 0.83 | 0.31 | <sup>16</sup> |
| A | eKC408 | ST - mrdA |  | 1.0715 | 0.1358 | 131 | 0.95 | 0.41 | <sup>16</sup> |
| A | eKC409 | VC - mrdA |  | 1.4799 | 0.1121 | 773 | 0.42 | 0.16 | <sup>16</sup> |
| A | eKC410 | YP - mrdA |  | 1.1447 | 0.1887 | 457 | 0.85 | 0.31 | <sup>16</sup> |
| A | eKC717 |  | 0.005% Arab. | 1.4657 | 0.2501 | 775 | 0.33 | 0.13 | <sup>15</sup> |
| A | eKC717 |  | 0.01% Arab. | 1.2809 | 0.1892 | 561 | 0.44 | 0.25 | <sup>15</sup> |
| A | eKC717 |  | 0.05% Arab. | 1.229 | 0.149 | 794 | 0.88 | 0.34 | <sup>15</sup> |
| A | eKC717 |  | 0.1% Arab. | 1.2239 | 0.2842 | 575 | 0.88 | 0.46 | <sup>15</sup> |
| A | eKC717 |  | 0.2% Arab. | 1.194 | 0.2329 | 239 | 0.90 | 0.34 | <sup>15</sup> |
| B | AV88 pAV20 G5 | WT – no repression | 100 ng/ml ATc | 1.1132 | 0.1044 | 812 | 0.93 | 0.31 | This work. |
| B | AV88 pAV20 G10 |  | 100 ng/ml ATc | 1.1284 | 0.1981 | 691 | 0.80 | 0.37 | This work. |
| B | AV88 pAV20 G14 |  | 100 ng/ml ATc | 1.6528 | 0.308 | 531 | 0.23 | 0.21 | This work. |
| B | AV88 pAV20 G20 |  | 100 ng/ml ATc | 1.9188 | 0.278 | 451 | 0.15 | 0.09 | This work. |
| B | AV88 pcrRNA G10 |  | 100 ng/ml ATc | 1.1355 | 0.1052 | 1103 | 0.89 | 0.44 | This work. |
| B | AV88 pcrRNA G11 |  | 100 ng/ml ATc | 1.2541 | 0.1008 | 878 | 0.56 | 0.33 | This work. |
| B | AV88 pcrRNA G14 |  | 100 ng/ml ATc | 1.2938 | 0.1323 | 342 | 0.56 | 0.34 | This work. |
| B | AV88 pcrRNA G20 |  | 100 ng/ml ATc | 1.9238 | 0.3106 | 409 | 0.08 | 0.07 | This work. |

**Table S2.** – *B. subtilis* strains used in this study

| Strain | Relevant genotype | Source |
| --- | --- | --- |
| bMD277 | *amyE::Pxyl-mreBCD, minCD (B. megaterium)::erm* | This work |
| bMD465 | *amyE::Pxyl-mreBCD, minCD (B. megaterium)::erm, mreBCD, minC,D::spec (B. megaterium)* | This work |
| bMK355 | *amyE::Pxyl-mreBCD::erm* | This work |
| bMD543 | *amyE::Pxyl-mreBCD::erm ΔmreBCD, ΔminCD::cat* | This work |
| bMD545 | *amyE::Pxyl-mreBCD::erm ΔmreBCD, PmreB-minCD::spec* | This work |
| bMD599 | *ΔponA::kan* | This work |
| bMK005 | *ΔponA::cat* | 17 |
| bMD554 | *yhdG::Phyperspank-ponA::cat* | This work |
| bMD586 | *yhdG::Phyperspank-ponA::cat ΔponA::kan* | This work |
| bMD594 | *yhdG::Pspank-ponA::cat* | This work |
| bMD598 | *yhdG::Pspank-ponA::cat ΔponA::kan* | This work |
| bMD619 | *amyE::Pxyl-mreBCD::erm ΔmreBCD, PmreB-minCD::spec yhdG::Pspank-ponA::cat* | This work |
| bMD620 | *amyE::Pxyl-mreBCD::erm ΔmreBCD, PmreB-minCD::spec yhdG::Pspank-ponA::cat ΔponA::kan* | This work |
| bMD622 | *amyE::Pxyl-mreBCD::erm yhdG::Pspank-ponA::cat ΔponA::kan* | This work |
| bMD556 | *yhdG::Phyperspank-rodA::cat* | This work |
| bMD580 | *yhdG::Phyperspank-rodA::cat ΔrodA::kan* | This work |
| bMD592 | *Pxyl-rodA::erm* | This work |
| bDR2487 | *ΔpbpH::spec* | 18 |
| bMD627 | *Pxyl-rodA::erm ΔpbpH::spec* | This work |
| bMD563 | *yhdG::Phyperspank-pbpA::cat* | This work |
| bRB776 | *yhdG::Pspank-pbpH::phleo ΔpbpH::spec ΔpbpA::erm* | 18 |
| bMD573 | *yhdG::Phyperspank-pbpA::cat ΔpbpH::spec* | This work |
| bMD574 | *yhdG::Phyperspank-pbpA::cat ΔpbpA::erm ΔpbpH::spec* | This work |
| bMD590 | *Pxyl-pbpA::erm* | This work |
| bMD597 | *Pxyl-pbpA::erm ΔpbpH::spec* | This work |
| bMD557 | *amyE::Pxyl-mreBCD::erm yhdG::Phyperspank-rodA::cat* | This work |
| bMD571 | *amyE::Pxyl-mreBCD::erm yhdG::Phyperspank-pbpA::cat* | This work |
| bRB773 | *yhdG::Pspank-pbpA::phleo ΔpbpH::spec ΔpbpA::cat* | 18 |
| bMD631 | *yhdG::Pspank-pbpA::phleo ΔpbpH::spec ΔpbpA::cat Pxyl-rodA::erm* | This work |
| bYS201 | *HaloTag-11aa-pbpA::cat* | This work |
| bMK385 | *amyE::Pxyl-mreBCD::erm ΔmreBCD, PmreB-minCD::spec HaloTag-11aa-pbpA::cat* | This work |
| bAM268 | *ΔpbpF ΔpbpG ΔpbpD ΔponA::kan* | 19 |
| bAM288 | *ΔpbpF ΔpbpG ΔpbpD ΔponA::kan amyE::Phyperspank-rodA-His10::spec* | 19 |
| bMD88 | *Pxyl-mreC::erm* | 12 |
| bYS09 | *mreB-mNeonGreen* | 20 |
| bYS19 | *mreB-SW-msfGFP* | This work |
| bYS977 | *amyE::Pxyl-mreB-SW-msfGFP,mreC,mreD::erm* | This work |
| bYS979 | *amyE::Pxyl-mreB-SW-msfGFP,mreC,mreD::erm, ΔmreBCD, ΔminCD::cat* | This work |
| bYS981 | *amyE::Pxyl-mreB-SW-msfGFP,mreC,mreD::erm, ΔmreBCD, PmreB-minC,D::spec* | This work |

**Table S3.** – *E. coli* strains used in this study

| Strain | Relevant genotype | Source |
| --- | --- | --- |
| MG1655 | Wild-type *E. coli* | CGSC #6300 |
| NO50 | *mreB-msfGFP sw* | 13 |
| NO50 - A20S | *NO50 with mreB(A20S)* | 13 |
| NO50 - L322Q | *NO50 with mreB(L322Q)* | 13 |
| NO50 - ΔA53 | *NO50 with mreB(ΔA53)* | 13 |
| NO50 - S185F | *NO50 with mreB(S185F)* | 13 |
| NO50 - F84V | *NO50 with mreB(F84V)* | 13 |
| NO50 - R74C | *NO50 with mreB(R74C)* | 13 |
| NO50 - S14A | *NO50 with mreB(S14A)* | 14 |
| RM478 | *ΔrodZ, mreB(S14A)-msfGFP* | 14 |
| KC407 | *ΔmrdA csrA::frt, mreB::msfGFP-mreB" + pww308 Ec mrdA* | 16 |
| KC408 | *ΔmrdA csrA::frt, mreB::msfGFP-mreB" + pww308 St mrdA* | 16 |
| KC409 | *ΔmrdA csrA::frt, mreB::msfGFP-mreB" + pww308 Vc mrdA* | 16 |
| KC410 | *ΔmrdA csrA::frt, mreB::msfGFP-mreB" + pww308 Yp mrdA* | 16 |
| NO34 | *csrD::kan, mreB::msfGFP-mreB* | 13 |
| KC507 | *csrD::kan, mreB::msfGFP-mreB<sup>E276D</sup>* | 15 |
| KC717 | *csrD::kan, mreB::msfGFP-mreB,ProdZ◊(frt araC PBAD)* | 15 |
| KC967 | *csrD::kan, mreB::msfGFP-mreB'-D83E* | 15 |
| KC968 | *csrD::kan, mreB::msfGFP-mreB'-R124C* | 15 |
| KC969 | *csrD::kan, mreB::msfGFP-mreB'-A174T* | 15 |
| AV88 | *186::P<sub>Tet</sub>-dCas9, mreB::msfGFP-mreB* | This work |
| AV88 pAV20 G5 | *AV88 with pAV20 G5* | This work |
| AV88 pAV20 G10 | *AV88 with pAV20 G10* | This work |
| AV88 pAV20 G14 | *AV88 with pAV20 G14* | This work |
| AV88 pAV20 G20 | *AV88 with pAV20 G20* | This work |
| AV88 pcrRNA G10 | *AV88 with pcrRNA G10* | This work |
| AV88 pcrRNA G11 | *AV88 with pcrRNA G11* | This work |
| AV88 pcrRNA G14 | *AV88 with pcrRNA G14* | This work |
| AV88 pcrRNA G20 | *AV88 with pcrRNA G20* | This work |

**Table S4.** – Plasmids used in this study

|  |  |  |
| --- | --- | --- |
| *pAV20 G5* | *guide RNA is sgRNA pAV20 G5, 5 matching base pairs against GFP* | This work |
| *pAV20 G10* | *guide RNA is sgRNA pAV20 G10, 10 matching base pairs against GFP* | This work |
| *pAV20 G14* | *guide RNA is sgRNA pAV20 G14, 14 matching base pairs against GFP* | This work |
| *pAV20 G20* | *guide RNA is sgRNA pAV20 G20, 20 matching base pairs against GFP* | This work |
| *pcrRNA G10* | *guide RNA is native crRNA, 10 matching base pairs against GFP* | This work |
| *pcrRNA G11* | *guide RNA is native crRNA, 11 matching base pairs against GFP* | This work |
| *pcrRNA G14* | *guide RNA is native crRNA, 14 matching base pairs against GFP* | This work |
| *pcrRNA G20* | *guide RNA is native crRNA, 20 matching base pairs against GFP* | This work |

**Table S5.**
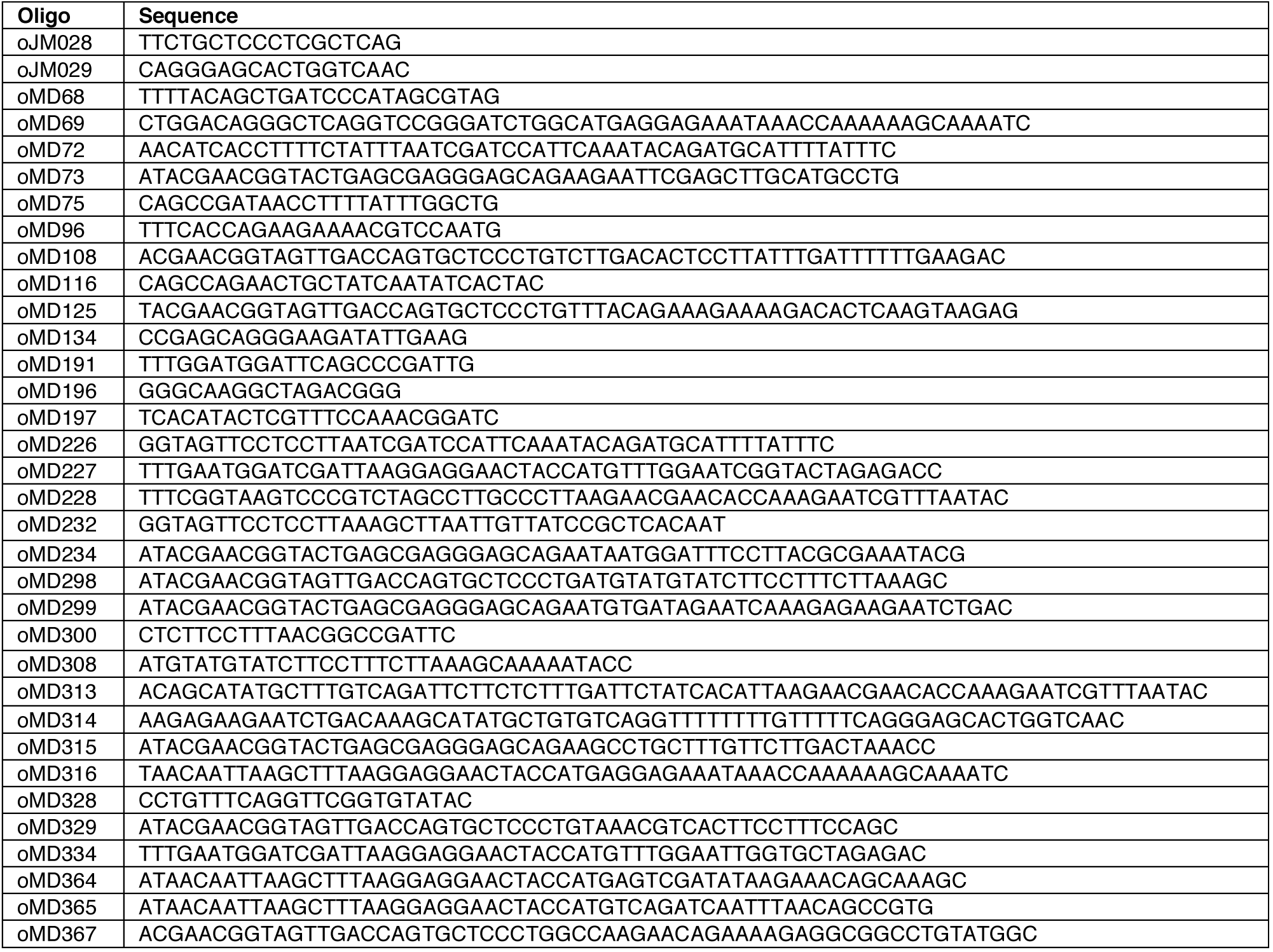
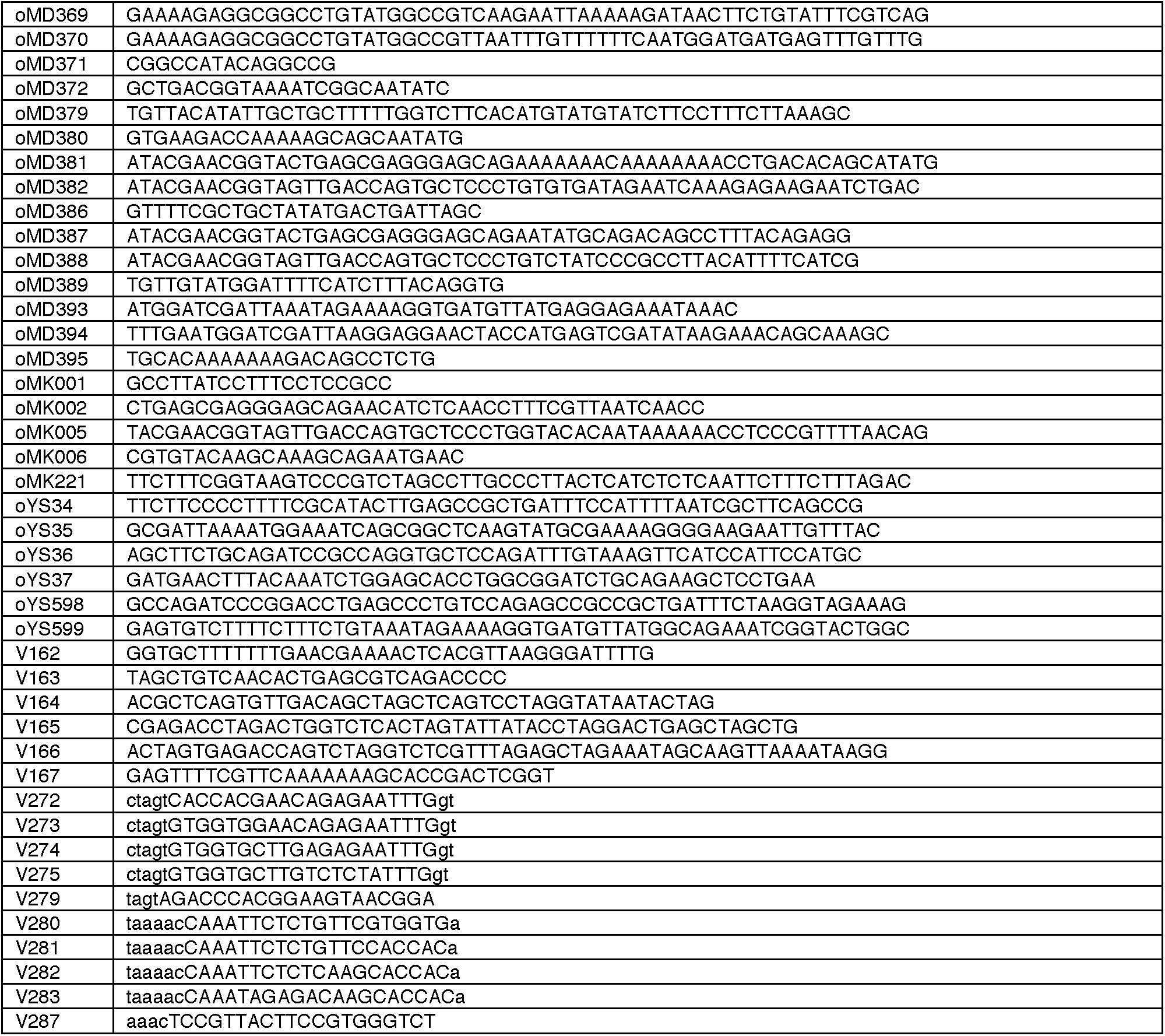
– Oligonucleotides used in this study

## References

1. Vadia, S. & Levin, P. A. Growth rate and cell size: a re-examination of the growth law. Curr Opin Microbiol 24, 96–103 (2015).

2. Sharpe, M. E., Hauser, P. M., Sharpe, R. G. & Errington, J. Bacillus subtilis cell cycle as studied by fluorescence microscopy: constancy of cell length at initiation of DNA replication and evidence for active nucleoid partitioning. J Bacteriol 180, 547–555 (1998).

3. Vollmer, W., Blanot, D. & de Pedro, M. A. Peptidoglycan structure and architecture. FEMS Microbiol Rev 32, 149–167 (2008).

4. Lebar, M. D. et al. Reconstitution of Peptidoglycan Cross-Linking Leads to Improved Fluorescent Probes of Cell Wall Synthesis. J Am Chem Soc 136, 10874–10877 (2014).

5. Banzhaf, M. et al. Cooperativity of peptidoglycan synthases active in bacterial cell elongation. Mol Microbiol 85, 179–194 (2012).

6. Meeske, A. J. et al. SEDS proteins are a widespread family of bacterial cell wall polymerases. Nature 537, 634–638 (2016).

7. Jones, L. J., Carballido-López, R. & Errington, J. Control of cell shape in bacteria: helical, actin-like filaments in Bacillus subtilis. Cell 104, 913–922 (2001).

8. van den Ent, F., Amos, L. & Löwe, J. Bacterial ancestry of actin and tubulin. Curr Opin Microbiol 4, 634–638 (2001).

9. van den Ent, F., Izoré, T., Bharat, T. A., Johnson, C. M. & Lowe, J. Bacterial actin MreB forms antiparallel double filaments. Elife 3, e02634 (2014).

10. Salje, J., van den Ent, F., de Boer, P. & Lowe, J. Direct membrane binding by bacterial actin MreB. Mol Cell 43, 478–487 (2011).

11. Hussain, S. et al. MreB filaments align along greatest principal membrane curvature to orient cell wall synthesis. Elife 7, 1239 (2018).

12. Garner, E. C. et al. Coupled, circumferential motions of the cell wall synthesis machinery and MreB filaments in B. subtilis. Science 333, 222–225 (2011).

13. Domínguez-Escobar, J. et al. Processive Movement of MreB-Associated Cell Wall Biosynthetic Complexes in Bacteria. Science 333, 225–228 (2011).

14. Cho, H. et al. Bacterial cell wall biogenesis is mediated by SEDS and PBP polymerase families functioning semi-autonomously. Nat Microbiol 1, 16172 (2016).

15. van Teeffelen, S. et al. The bacterial actin MreB rotates, and rotation depends on cell-wall assembly. Proc Natl Acad Sci USA 108, 15822–15827 (2011).

16. Turner, R. D., Mesnage, S., Hobbs, J. K. & Foster, S. J. Molecular imaging of glycan chains couples cell-wall polysaccharide architecture to bacterial cell morphology. Nat Comms 9, 1263 (2018).

17. McPherson, D. C. & Popham, D. L. Peptidoglycan synthesis in the absence of class A penicillin-binding proteins in Bacillus subtilis. J Bacteriol 185, 1423–1431 (2003).

18. Ursell, T. S. et al. Rod-like bacterial shape is maintained by feedback between cell curvature and cytoskeletal localization. Proc Natl Acad Sci USA 111, E1025–34 (2014).

19. Ouzounov, N. et al. MreB Orientation Correlates with Cell Diameter in Escherichia coli. Biophys J 111, 1035–1043 (2016).

20. Shi, H., Bratton, B. P., Gitai, Z. & Huang, K. C. How to Build a Bacterial Cell: MreB as the Foreman of E. coli Construction. Cell 172, 1294–1305 (2018).

21. Wang, S. & Wingreen, N. S. Cell shape can mediate the spatial organization of the bacterial cytoskeleton. Biophys J 104, 541–552 (2013).

22. Tropini, C. et al. Principles of Bacterial Cell-Size Determination Revealed by Cell-Wall Synthesis Perturbations. Cell Reports 9, 1520–1527 (2014).

23. Shi, H. et al. Deep Phenotypic Mapping of Bacterial Cytoskeletal Mutants Reveals Physiological Robustness to Cell Size. Curr Biol 27, 3419–3429.e4 (2017).

24. Colavin, A., Shi, H. & Huang, K. C. RodZ modulates geometric localization of the bacterial actin MreB to regulate cell shape. Nat Comms 9, 1280 (2018).

25. Olshausen, P. V. et al. Superresolution Imaging of Dynamic MreB Filaments in B. subtilis—A Multiple-Motor-Driven Transport? Biophys J 105, 1171–1181 (2013).

26. Shi, H., Bratton, B. P., Gitai, Z. & Huang, K. C. How to Build a Bacterial Cell: MreB as the Foreman of E. coli Construction. Cell 172, 1294–1305 (2018).

27. Harris, L. K., Dye, N. A. & Theriot, J. A. A Caulobacter MreB mutant with irregular cell shape exhibits compen& - PubMed - NCBI. Mol Microbiol 94, 988–1005 (2014).

28. Vigouroux, A., Oldewurtel, E., Cui, L., Bikard, D. & van Teeffelen, S. Tuning dCas9’s ability to block transcription enables robust, noiseless knockdown of bacterial genes. Molecular Systems Biology 14, e7899 (2018).

29. Henriques, A. O., Glaser, P., Piggot, P. J. & Moran, C. P., Jr. Control of cell shape and elongation by the rodA gene in Bacillus subtilis. Mol Microbiol 28, 235–247 (1998).

30. Fraipont, C. et al. The integral membrane FtsW protein and peptidoglycan synthase PBP3 form a subcomplex in Escherichia coli. Microbiology (Reading, Engl) 157, 251–259 (2010).

31. Harris, L. K. & Theriot, J. A. Relative Rates of Surface and Volume Synthesis Set Bacterial Cell Size. Cell 165, 1479–1492 (2016).

32. Taheri-Araghi, S. et al. Cell-Size Control and Homeostasis in Bacteria. Curr Biol 27, 1392 (2017).

33. Zheng, H. et al. Interrogating the Escherichia coli cell cycle by cell dimension perturbations. Proc Natl Acad Sci USA 113, 15000–15005 (2016).

34. Murray, T., Popham, D. L. & Setlow, P. Bacillus subtilis cells lacking penicillin-binding protein 1 require increased levels of divalent cations for growth. (1998).

35. Popham, D. L. & Setlow, P. Phenotypes of Bacillus subtilis mutants lacking multiple class A high-molecular-weight penicillin-binding proteins. J Bacteriol 178, 2079–2085 (1996).

36. Emami, K. et al. RodA as the missing glycosyltransferase in Bacillus subtilis and antibiotic discovery for the peptidoglycan polymerase pathway. Nat Microbiol 2, 16253 (2017).

37. Billaudeau, C. et al. Contrasting mechanisms of growth in two model rod-shaped bacteria. Nat Comms 8, 15370 (2017).

38. Grimm, J. B. et al. A general method to improve fluorophores for live-cell and single-molecule microscopy. Nat Meth 12, 244–250 (2015).

39. Oldenbourg, R. Polarized light microscopy: principles and practice. Cold Spring Harb Protoc 2013, (2013).

40. Probine, M. C. & Preston, R. D. Cell Growth and the Structure and Mechanical Properties of the Wall in Internodal Cells of Nitella opacaI. Wall structure and growth. J Exp Bot 12, 261–282 (1961).

41. Abraham, Y. & Elbaum, R. Quantification of microfibril angle in secondary cell walls at subcellular resolution by means of polarized light microscopy. New Phytol 197, 1012–1019 (2012).

42. Baskin, T. I. Anisotropic expansion of the plant cell wall. Annu Rev Cell Dev Biol 21, 203–222 (2005).

43. Inouïï, S. Polarization Microscopy. Current protocols in cell biology 1, (John Wiley & Sons, Inc., 2001).

44. Verwer, R. W., Beachey, E. H., Keck, W., Stoub, A. M. & Poldermans, J. E. Oriented fragmentation of Escherichia coli sacculi by sonication. J Bacteriol 141, 327–332 (1980).

45. Gan, L., Chen, S. & Jensen, G. J. Molecular organization of Gram-negative peptidoglycan. Proc Natl Acad Sci USA 105, 18953–18957 (2008).

46. Yao, X., Jericho, M., Pink, D. & Beveridge, T. Thickness and elasticity of gram-negative murein sacculi measured by atomic force microscopy. J Bacteriol (1999).

47. Cui, L. et al. A CRISPRi screen in E. coli reveals sequence-specific toxicity of dCas9. Nat Comms 9, 1912 (2018).

48. Bratton, B. P., Shaevitz, J. W., Gitai, Z. & Morgenstein, R. M. MreB polymers and curvature localization are enhanced by RodZ and predict E. coli’s cylindrical uniformity. - PubMed - NCBI. Nat Comms 9, 12510 (2018).

49. Typas, A., Banzhaf, M., Gross, C. A. & Vollmer, W. From the regulation of peptidoglycan synthesis to bacterial growth and morphology. Nat Rev Microbiol 10, 123–136 (2011).

50. Essential nature of the mreC determinant of Bacillus subtilis. J Bacteriol 185, 4490–4498 (2003).

51. Leaver, M. & Errington, J. Roles for MreC and MreD proteins in helical growth of the cylindrical cell wall in Bacillus subtilis. Mol Microbiol 57, 1196–1209 (2005).

52. Fisher, I. C., Shapiro, L. & Theriot, J. A. Mutations in the nucleotide binding pocket of MreB can alter cell curvature and polar morphology in Caulobacter. Mol Microbiol 81, 368–394 (2011).

53. Land, A. D. The requirement for pneumococcal MreC and MreD is relieved by inactivation of the gene encoding PBP1a. J Bacteriol 193, 4166–4179 (2011).

54. Kawai, Y., Daniel, R. A. & Errington, J. Regulation of cell wall morphogenesis in Bacillus subtilis by recruitment of PBP1 to the MreB helix. Mol Microbiol 71, 1131–1144 (2009).

55. Lai, G. C., Cho, H. & Bernhardt, T. G. The mecillinam resistome reveals a role for peptidoglycan endopeptidases in stimulating cell wall synthesis in Escherichia coli. PLoS Genet 13, e1006934 (2017).

56. Lee, T. K., Meng, K., Shi, H. & Huang, K. C. Single-molecule imaging reveals modulation of cell wall synthesis dynamics in live bacterial cells. Nat Comms 7, 13170 (2016).

## Supplemental References

1. Grimm, J. B. et al. A general method to improve fluorophores for live-cell and single-molecule microscopy. Nat Meth 12, 244–250 (2015).

2. Ursell, T. et al. Rapid, precise quantification of bacterial cellular dimensions across a genomic-scale knockout library. BMC Biology 15, 17 (2017).

3. Vadia, S. & Levin, P. A. Growth rate and cell size: a re-examination of the growth law. Curr Opin Microbiol 24, 96–103 (2015).

4. Ursell, T. S. et al. Rod-like bacterial shape is maintained by feedback between cell curvature and cytoskeletal localization. Proc Natl Acad Sci USA 111, E1025–34 (2014).

5. Wiseman, P. W. Image correlation spectroscopy: mapping correlations in space, time, and reciprocal space. - PubMed - NCBI. Fluorescence Fluctuation Spectroscopy (FFS), Part A 518, 245–267 (2013).

6. Sharpe, M. E., Hauser, P. M., Sharpe, R. G. & Errington, J. Bacillus subtilis cell cycle as studied by fluorescence microscopy: constancy of cell length at initiation of DNA replication and evidence for active nucleoid partitioning. J Bacteriol 180, 547–555 (1998).

7. Vollmer, W., Blanot, D. & de Pedro, M. A. Peptidoglycan structure and architecture. FEMS Microbiol Rev 32, 149–167 (2008).

8. Kner, P., Chhun, B. B., Griffis, E. R., Winoto, L. & Gustafsson, M. G. L. Super-resolution video microscopy of live cells by structured illumination. Nat Meth 6, 339–342 (2009).

9. Vigouroux, A., Oldewurtel, E., Cui, L., Bikard, D. & van Teeffelen, S. Tuning dCas9’s ability to block transcription enables robust, noiseless knockdown of bacterial genes. Molecular Systems Biology 14, e7899 (2018).

10. Kall, L., Storey, J. D. & Noble, W. S. Non-parametric estimation of posterior error probabilities associated with peptides identified by tandem mass spectrometry. Bioinformatics 24, i42–i48 (2008).

11. Gibson, D. G. et al. Enzymatic assembly of DNA molecules up to several hundred kilobases. Nat Meth 6, 343–U41 (2009).

12. Schirner, K. et al. Lipid-linked cell wall precursors regulate membrane association of bacterial actin MreB. Nat. Chem. Biol. 11, 38–45 (2015).

13. Ouzounov, N. et al. MreB Orientation Correlates with Cell Diameter in Escherichia coli. Biophys J 111, 1035–1043 (2016).

14. Morgenstein, R. M. et al. RodZ links MreB to cell wall synthesis to mediate MreB rotation and robust morphogenesis. Proc Natl Acad Sci USA 112, 12510–12515 (2015).

15. Colavin, A., Shi, H. & Huang, K. C. RodZ modulates geometric localization of the bacterial actin MreB to regulate cell shape. Nat Comms 9, 1280 (2018).

16. Tropini, C. et al. Principles of Bacterial Cell-Size Determination Revealed by Cell-Wall Synthesis Perturbations. Cell Reports 9, 1520–1527 (2014).

17. Cho, H. et al. Bacterial cell wall biogenesis is mediated by SEDS and PBP polymerase families functioning semi-autonomously. Nat Microbiol 1, 16172 (2016).

18. Garner, E. C. et al. Coupled, circumferential motions of the cell wall synthesis machinery and MreB filaments in B. subtilis. Science 333, 222–225 (2011).

19. Meeske, A. J. et al. SEDS proteins are a widespread family of bacterial cell wall polymerases. Nature 537, 634–638 (2016).

20. Hussain, S. et al. MreB filaments align along greatest principal membrane curvature to orient cell wall synthesis. Elife 7, 1239 (2018).

